# A comparative analysis of microglial inducible Cre lines

**DOI:** 10.1101/2023.01.09.523268

**Authors:** Travis E. Faust, Philip A. Feinberg, Ciara O’Connor, Riki Kawaguchi, Andrew Chan, Haley Strasburger, Takahiro Masuda, Lukas Amann, Klaus-Peter Knobeloch, Marco Prinz, Anne Schaefer, Dorothy P. Schafer

**Author notes:** authors contributed equally (Travis E. Faust, Philip A. Feinberg).

## Abstract

Cre/LoxP technology has revolutionized genetic studies and allowed for spatial and temporal control of gene expression in specific cell types. The field of microglial biology has particularly benefited from this technology as microglia have historically been difficult to transduce with virus or electroporation methods for gene delivery. Here, we interrogate four of the most widely available microglial inducible Cre lines. We demonstrate varying degrees of recombination efficiency and spontaneous recombination, depending on the Cre line and loxP distance. We also establish best practice guidelines and protocols to measure recombination efficiency in microglia, which could be extended to other cell types. There is increasing evidence that microglia are key regulators of neural circuit structure and function. Microglia are also major drivers of a broad range of neurological diseases. Thus, reliable manipulation of their function in vivo is of utmost importance. Identifying caveats and benefits of all tools and implementing the most rigorous protocols are crucial to the growth of the field of microglial biology and the development of microglia-based therapeutics.

## Introduction

Microglia are resident central nervous system (CNS) macrophages now appreciated to play key roles in regulation of synaptic connectivity and modulation of neural activity (Prinz et al., 2019; Wu et al., 2015). Microglia are also increasingly recognized as important cellular drivers of brain diseases, ranging from neurodegenerative diseases such as Alzheimer Disease (AD) to neuropsychiatric disorders such as schizophrenia (Biber et al., 2016; Salter and Stevens, 2017). This includes studies that have identified many microglia-enriched genes as key risk factors (Hansen et al., 2018), which are likely contributing to the underlying etiology of disease. Thus, methods to study microglial gene function in a cell type-specific way is of utmost importance. Until very recently, microglia were not easily targeted with viruses for genetic manipulation (Lin et al., 2022). In addition, electroporation for gene delivery has been ineffective and Tet-On/Off systems require experiments to be carried out in the presence of immunosuppressive drugs. As a result, the Cre/Lox system has been the primary method to genetically manipulate microglia *in vivo* and a number of microglial Cre lines have been made in recent years. Here, we provide a comprehensive comparison of some of the most widely available tamoxifen-inducible microglial Cre lines and a guide for best practices to ensure efficient and specific recombination.

When the Cre protein is fused to a mutated estrogen-binding receptor domain (ER), the fusion CreER protein is retained to the cytoplasm and prevents the Cre-mediated recombination of LoxP sequences in the genomic DNA (Feil et al., 1996). Once the CreER binds 4- hydroxytamoxifen (4-OHT), the active metabolite of tamoxifen (TAM), the CreER translocates to the nucleus and recombination can occur. Recombination efficiency depends on several factors, including the level of CreER expression, the amount of TAM that enters the cell, the accessibility of the LoxP locus, and the inter-loxP distance of the floxed alleles (Glaser et al., 2005). These variables necessitate validation of Cre/LoxP recombination in each experimental paradigm (McLellan et al., 2017). The most widely used CreER lines to target microglia have leveraged the promoter for the gene encoding the fractalkine receptor (CX3CR1) (Goldmann et al., 2013; Parkhurst et al., 2013; Yona et al., 2013). Compared to other recently generated lines such as *Sall1*^CreER^ (Chappell-Maor et al., 2020) or *Cd11b^Cre^* (Shi et al., 2018), there is less recombination in other brain or immune cell types. *Cx3cr1* is also expressed by other peripheral immune cells, such as circulating, bone-marrow derived monocytes. However, the majority of these peripheral cells are short-lived and turnover within 4 weeks due to ongoing, lifelong hematopoiesis (Goldmann et al., 2013; Parkhurst et al., 2013; Yona et al., 2013). Thus, 4 weeks after TAM administration, recombination is largely restricted to microglia and other long-lived macrophages (e.g., perivascular macrophages) in *Cx3cr1^CreER^* mice (Goldmann et al., 2016). While these mice have led to significant advancements in the field as the first more-specific microglial genetic tool, there are some limitations. First, these are knock-in lines in which *CreER* was knocked into the *Cx3cr1* locus in a way that disrupts endogenous *Cx3cr1* gene expression. While most groups use them as heterozygotes to avoid knocking out *Cx3cr1*, these heterozygous mice still lack a copy of the *Cx3cr1* gene. In addition, the need for peripheral macrophage turnover after 4 weeks following TAM administration limits interpretation of studies during the first few weeks of postnatal development. There are also specific CX3CR1+ populations of longer-lived tissue resident macrophages that, similar to microglia, still remain recombined after 4 weeks following TAM administration, such as those in the leptomeninges and the perivascular space (Goldmann et al., 2016). In certain cases, it has also been shown that *Cx3cr1^CreER^* can induce LoxP-recombination in microglia in the absence of TAM (Fonseca et al., 2017; Stowell et al., 2019; Van Hove et al., 2020). Also, TAM administration in early postnatal *Cx3cr1^CreER^* pups can induce an interferon responsive population of microglia later in development (Sahasrabuddhe and Ghosh, 2022). These limitations have led to the recent development of new inducible Cre lines that more specifically target microglia vs. other macrophage populations, including *Tmem119^CreER^* (Kaiser and Feng, 2019), *Hexb^CreER^* (Masuda et al., 2020), and *P2ry12^CreER^* mice (McKinsey et al., 2020). Another recent development is the generation of a non-inducible, highly specific microglial split-Cre line, for experiments where temporal regulation of gene recombination is not required (Kim et al., 2021).

The effectiveness of CreER lines depends on several factors such as: 1) the cell-type specificity of the CreER expression pattern, 2) the expression level and ability to efficiently induce LoxP-recombination, 3) a lack of spontaneous recombination in the absence of TAM, and 4) a lack of off-target effects in both the presence and absence of TAM. Therefore, several quality control experiments should be performed. In the current study, we performed a systematic comparison of four of the most widely available inducible microglial CreER lines: *Tmem119^CreER^*, *Hexb^CreER^*, and two different *Cx3cr1^CreER^* lines. Together, our results demonstrate that recombination efficiency varies across these CreER lines. Our data further support that recombination efficiency can be boosted by breeding the CreER to homozygosity for some lines and using floxed mouse lines with shorter inter- LoxP distances. Care should also be taken as some CreER lines induce a higher degree of spontaneous recombination. We finally provide guidelines, quality control measures, and methods for future use of the Cre/LoxP system in microglia, which can be extended to other cell types.

## Results

### Recombination efficiency varies across microglial CreER lines

We initially compared recombination across the *Cx3cr1^YFP-CreER (Litt)^* (JAX #021160) (Parkhurst et al., 2013), *Cx3cr1^CreER (Jung)^* (JAX #020940) (Yona et al., 2013), *Tmem119^CreER^* (JAX #031820) (Kaiser and Feng, 2019), and *Hexb^CreER^* (Masuda et al., 2020) mouse lines using the *Rosa26^mTmG^* reporter line (Muzumdar et al., 2007). This reporter line expresses LoxP- [membrane tdTomato (mTomato)]-STOP-LoxP-[membrane GFP (mGFP)] under the *Rosa26* promoter (Fig. 1A). For these initial experiments we used mice heterozygous for the *Rosa26^mTmG^* allele such that only one allele will be recombined for a given cell. At baseline, all cells express mTomato, but in cells that undergo Cre/LoxP recombination, the mTomato and stop codon are removed, leading to mGFP expression. Mice were injected with TAM daily at postnatal day 28 (P28) through P31. At P56, half of each brain was used for fluorescence-activated cell sorting (FACS) and the other half of the brain was used for immunofluorescence microscopy (Fig. 1B). For FACS analyses, brains were mechanically dissociated, subjected to Percoll gradient, and immunostained for CD11b and CD45. Live microglia (DAPI^Negative^) were sorted by FACS as CD11b^High^ CD45^Mid^ (Fig. S1A). We then assessed recombination by evaluating the percentage of microglia that were unrecombined (mTomato+) vs. recombined (mGFP+) across the CreER lines (Fig. 1B; Fig. S1B). The percentage of recombined mGFP^+^ microglia was highest in heterozygous P56 *Cx3cr1^YFP-CreER/+ (Litt)^* (98.33% ± 0.32%) and *Cx3cr1^CreER/+ (Jung)^* (92.53% ± 2.08%) mice which had nearly 100% of cells recombined (Fig. 1C- E). In comparison, using the same TAM paradigm, recombination was less efficient in P56 *Tmem119^CreER/+^* (49.44% ± 3.47%) and *Hexb^CreER/+^* (33.16% ± 2.52%) mice (Fig. 1F-K).

**Figure 1.**
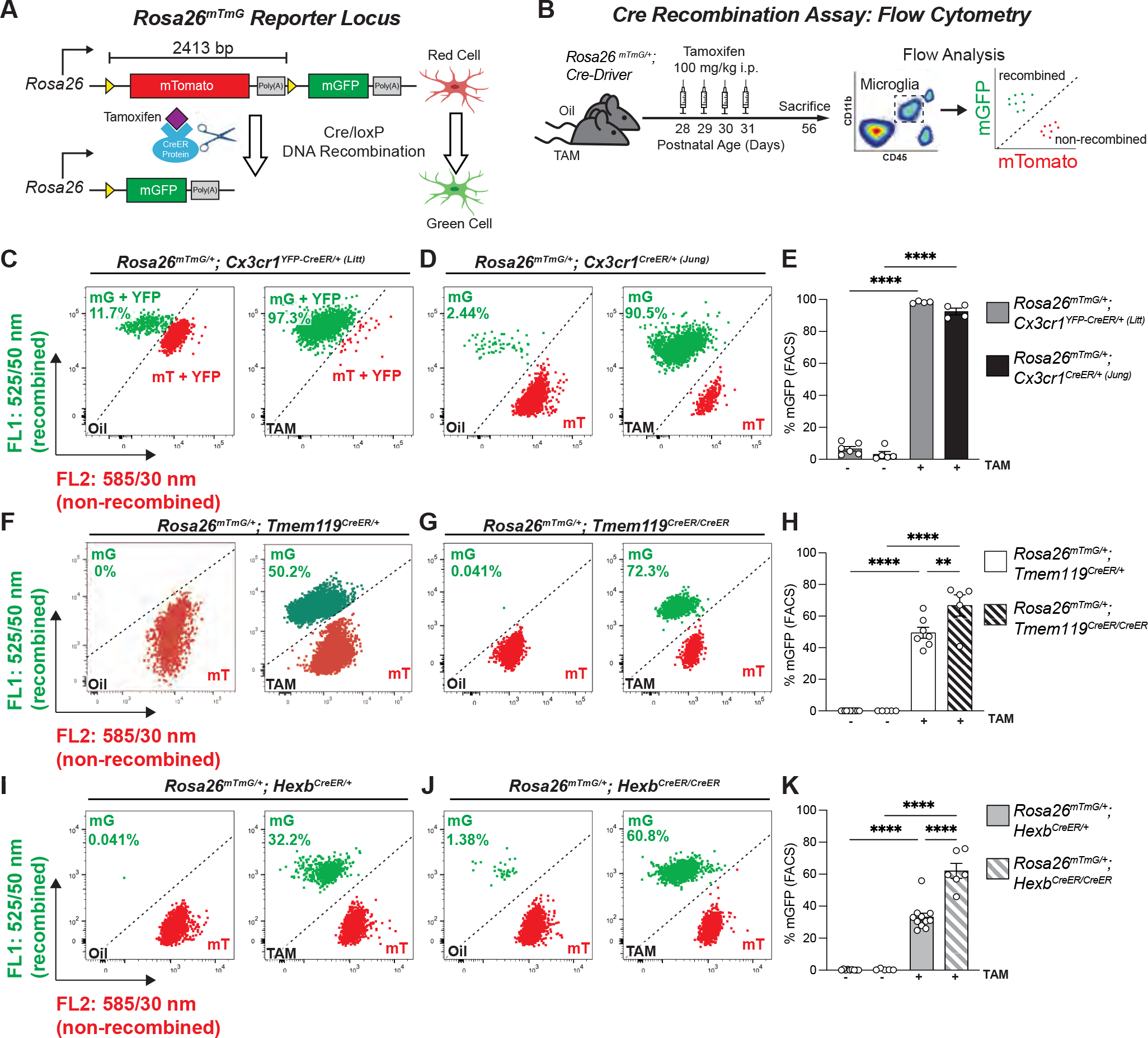
Recombination efficiency varies across microglial CreER lines. **A** Diagram of the *Rosa26^mTmG^* allele and corresponding cellular fluorescence before and after Cre/loxP DNA recombination. LoxP sites are indicated by yellow triangles. **B** Diagram of experimental protocol used to assess tamoxifen (TAM) induced Cre/loxP recombination of *Rosa26^mTmG/+^* in microglia by flow cytometry. **C,D,F,G,I,J**Representative flow cytometry results show the percentage of mGFP^+^ (mG^+^) and mTomato^+^ (mT^+^) microglia from individual animals from each group. **E,H,K** Quantification of the percentage of recombined mGFP^+^ microglia in *Cx3cr1^CreER^* lines (E), *TMEM119^CreER^* lines (H), and Hexb^CreER^ lines (K). 2-way ANOVA with Sidak’s post hoc. (E, *Cx3cr1^YFP-CreER/+ (Litt)^* oil vs TAM, *n =* 6 oil, 4 TAM mice, *P <* 0.0001, *t* = 41.37, *df* = 15; *Cx3cr1^CreER/+ (Jung)^* oil vs TAM, *n =* 5 oil, 4 TAM mice, *P <* 0.0001, *t* = 38.77, *df* = 15) (H, *Tmem119^CreER/+^* oil vs TAM, *n =* 11 oil, 7 TAM mice, *P <* 0.0001, *t* = 13.39, *df* = 24; *Tmem119^CreER/CreER^* oil vs TAM, *n =* 5 oil, 5 TAM mice, *P <* 0.0001, *t* = 16.2, *df* = 24; *Tmem119^CreER/+^* TAM vs *Tmem119^CreER/CreER^* TAM, *n =* 7 *CreER/+*, 5 *CreER/CreER* mice, *P <* 0.0001, *t* = 3.869, *df* = 24) (K, *Hexb^CreER/+^* oil vs TAM, *n =* 9 oil, 11 TAM mice, *P <* 0.0001, *t* = 10.36, *df* = 27; *Hexb^CreER/CreER^* oil vs TAM, *n =* 5 oil, 6 TAM mice, *P <* 0.0001, *t* = 14.40, *df* = 27; *Hexb^CreER/+^* TAM vs *Hexb^CreER/CreER^* TAM, *n =* 11 *CreER/+*, 6 *CreER/CreER*, *P <* 0.0001, *t* = 8.063, *df* = 27.) All data are presented as mean ± SEM. See also Figures S1 and S2.

Immunofluorescence microscopy of fixed brain sections from the same animals confirmed these results (Fig. 2D and Fig. S2). This immunofluorescence method is particularly useful for determining if recombination is restricted to microglia vs. other CNS cell types in intact tissue and if recombination is evenly distributed across the tissue.

**Figure 2.**
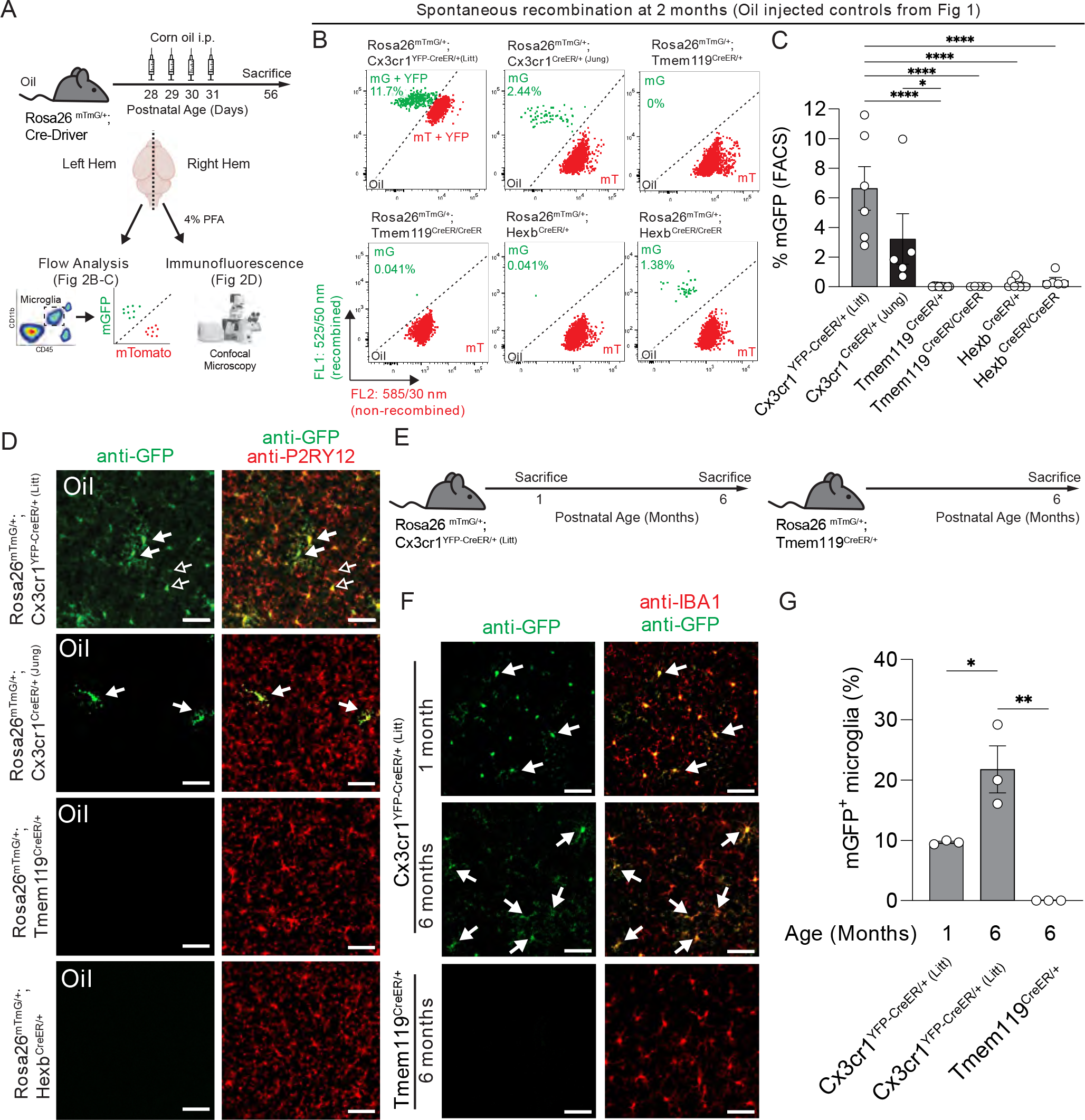
Spontaneous recombination occurs in the *Cx3cr1^CreER^* lines. **A** Diagram of experimental protocol used to assess spontaneous Cre/loxP recombination of *Rosa26^mTmG/+^* in microglia by flow cytometry and immunofluorescence. **B** Representative flow cytometry results show the percentage of recombined mGFP^+^ (mG) and mTomato^+^ (mT) microglia from individual animals from each group from Fig 1. **C** Quantification of the percentage of recombined mGFP^+^ microglia shows increased spontaneous recombination of the *Rosa26^mTmG^* allele in the *Cx3cr1^CreER^* lines compared to the *Tmem119^CreER^* and *Hexb^CreER^* lines (1-way ANOVA with Tukey’s post hoc; *Cx3cr1^YFP-CreER/+ (Litt)^* vs. *Tmem119^CreER/+^*, n = 6, 11 mice, *P <* 0.0001, *q* = 9.749, *df* = 35; *Cx3cr1^YFP-CreER/+ (Litt)^* vs. *Tmem119^CreER/CreER^*, n = 6, 5 mice, *P <* 0.0001, *q* = 8.171, *df* = 35; *Cx3cr1^YFP-CreER/+ (Litt)^* vs. *Hexb^CreER/+^*, n = 6, 9 mice, *P <* 0.0001, *q* = 9.114, *df* = 35; *Cx3cr1^YFP-CreER/+ (Litt)^* vs. *Hexb^CreER/CreER^*, n = 6, 5 mice, *P <* 0.0001, *q* = 7.675, *df* = 35; *Cx3cr1^CreER/+ (Jung)^* vs. *Tmem119^CreER/+^*, n = 5, 11 mice, *P <* 0.0368, *q* = 4.444, *df* = 35). **D** Representative immunofluorescent images of brain sections from right hemispheres of oil injected mice used for flow cytometry analysis in B-C. Sections were immunolabeled for anti- P2RY12 (red) to identify microglia and anti-GFP (green) to identify recombined cells. The number of recombined mGFP^+^ microglia (white arrows) matches the results observed by flow cytometry. In the *Cx3cr1^YFP-CreER/+ (Litt)^* line, the soma of unrecombined microglia are also immunolabeled by anti-GFP due to the constitutive expression YFP (open arrows), but can be distinguished from recombined mGFP^+^ microglia by fluorescence intensity and membrane labeling. Scale bars 50 µm. **E** Diagram of genotypes and ages used for assessment of spontaneous Cre/loxP recombination of *Rosa26^mTmG/+^* mice with no injection. **F** Representative immunofluorescent images of brain sections immunolabeled for anti-GFP (green) and anti-IBA1 (red) to identify recombined mGFP^+^ microglia (white arrows). **G** Quantification of the percentage of recombined mGFP^+^ microglia in the cortex shows increased recombination of *Rosa26^mTmG/+^* in 6 month old *Cx3cr1^YFP-CreER/+ (Litt)^* mice vs. 1 month old *Cx3cr1^YFP-CreER/+ (Litt)^* mice and 6 month old *Tmem119^CreER/+^* mice (1-way ANOVA with Tukey’s post hoc; *Cx3cr1^YFP-CreER/+ (Litt)^* 6 mo vs. 1 mo, *n* = 3 mice, *P* = 0.0012, *q* = 5.39*, df* = 6; *Cx3cr1^YFP-CreER/+ (Litt)^* 6 mo vs. *Tmem119^CreER/+^* 6 mo, *n* = 3 mice, *q* = 9.696*, df* = 6). Scale bars 50 µm. All data are presented as mean ± SEM.

Unlike the *Cx3cr1^CreER/+^* lines, the *Tmem119^CreER/+^* and *Hexb^CreER/+^* lines were engineered to insert a T2A/P2A-CreER protein cleavage sequence after the native gene sequence. As a result, use of mice homozygous for the CreER in these lines is a more viable strategy. We found that recombination of the *Rosa26*^mTmG^ allele was increased when crossed to homozygous *Tmem119^CreER/CreER^* (66.7% ± 6.74%) or *Hexb^CreER/CreER^* (62.12% ± 4.67%) mice (Fig. 1F-K).

Together, these results demonstrate highly efficient recombination in microglia with the *Cx3cr1*^CreER/+^ lines compared to *Tmem119^CreER/+^* and *Hexb^CreER/+^* lines. However, breeding *Tmem119^CreER/+^* and *Hexb^CreER/+^* lines to homozygosity is an effective strategy for increasing recombination efficiency in microglia.

### Assessment of spontaneous recombination

While the efficiency of TAM-dependent recombination was high in microglia in *Cx3cr1^YFP- CreER/+ (Litt)^* and *Cx3cr1^CreER/+ (Jung)^* mice, TAM-independent recombination events were regularly observed in the oil-treated controls for these lines (*Cx3cr1^YFP-CreER/+ (Litt)^* = 6.64% ± 1.48%; *Cx3cr1^CreER/+ (Jung)^* = 3.23% ± 1.71%) by FACS (Fig. 1C-Eand Fig. 2A-C). In contrast, spontaneous recombination was negligible by FACS in mice heterozygous or homozygous for *Tmem119*^CreER^ or *Hexb^CreER^* treated with oil (Fig. 1F-K and 2A-C). We observed similar levels of spontaneous recombination by immunofluorescence microscopy of fixed mouse brain sections from the same animals (Fig. 2A, D and Fig. S2). To further investigate this spontaneous recombination, we assessed *Cx3cr1^YFP-CreER/+ (Litt)^*; *Rosa26^mTmG^*^/+^ or *Tmem119^CreER/+^*; *Rosa26^mTmG/+^* mice with no TAM or oil injection (Fig 2e). In *Cx3cr1^CreER/+^*; *Rosa26^mTmG/+^* with no injection, we observed recombination (mGFP) in 9.7% ± 0.2% of cortical IBA1+ microglia at 1 month, which increased to 21.8% ± 3.9% of microglia by 6 months of age (Fig. 2F-G). These data are consistent with previous reports of TAM-independent, spontaneous recombination in *Cx3cr1^YFP-CreER/+ (Litt)^* mice (Fonseca et al., 2017; Stowell et al., 2019; Van Hove et al., 2020). In contrast, no spontaneous recombination events were observed in *Tmem119^CreER/+^*; *Rosa26^mTmG/+^* mice assessed at 6 months of age (Fig. 2F-G). To control for spontaneous recombination, appropriate oil-treated controls should be performed.

### Changes in gene expression in microglial CreER lines +/- tamoxifen treatment

Interestingly, in the process of analyzing spontaneous recombination in the *Cx3cr1^YFP- CreER/+ (Litt)^* mice, we noticed another important caveat. Similar to a recent publication (Sahasrabuddhe and Ghosh, 2022), we observed a downregulation of the homeostatic microglial marker P2RY12 in subpopulations of microglia when *Cx3cr1^YFP-CreER/+ (Litt)^* mice were injected with TAM neonatally and assessed at P28-P51 (Fig. S3). This phenotype was not observed in P28 *Tmem119^CreER/+^* mice treated with the same neonatal TAM paradigm or when *Cx3cr1^YFP-CreER/+ (Litt)^* mice were injected with TAM at 4 weeks of age (Fig. S2). These results suggest that this phenotype is likely not caused by the TAM itself, but a specific interaction between the TAM and the *Cx3cr1^YFP-CreER/+ (Litt)^* line during development. Also, it did not originate from a loss of one copy of *Cx3cr1* as Cx3cr1^EGFP/+^ mice, which also are heterozygous for CX3CR1, did not exhibit this phenotype when treated with TAM (Fig. S3A-B).

Based on the rather large differences in gene expression following TAM administration in *Cx3cr1^YFP-CreER/+ (Litt)^* neonates, we reasoned that there may be more subtle effects in gene expression in adults, too. We, therefore, more broadly compared gene expression across P56 CreER mice with and without TAM induction at P28-P31 by bulk RNA sequencing (RNA-Seq) (Fig. 3). Similar to other experiments (Figs. 1-2), all mice were injected daily with TAM or oil from P28-P31. Subsequently, microglia were acutely isolated from the brains and purified by FACS at P56 (Fig. 3A). Comparison of cell-type specific transcripts confirmed the enrichment of microglia (*Csf1r*, *P2ry12*) over other brain cell types (Fig. 3B). In addition, no major differences in gene expression were observed between CreER lines and TAM only a had minimal effect on gene expression (Fig. 3B-F). However, when we assessed gene expression of individual genes whose promoters are used to drive CreER expression across the different mouse lines, we found some significant differences. *Cx3cr1* mRNA was reduced by nearly 50% in *Cx3cr1^YFP-^ CreER/+ (Litt)* and Cx3cr1*CreER/+ (Jung)* microglia as compared to Hexb*CreER/CreER* or Tmem119*CreER/+* microglia (Fig. 3G). This was result was expected since the mice were made using a knock-in strategy in the *Cx3cr1* locus, which disrupts native gene expression (Parkhurst et al., 2013; Yona et al., 2013). Also, despite preserving the full coding region by inserting *P2A/T2A-CreER* constructs between the last exon and the stop codon in *Tmem119^CreER^* and *Hexb ^CreER^* mice, endogenous mRNA expression was still affected. *Tmem119* mRNA was reduced by ∼33% in *Tmem119*^CreER/+^ microglia as compared to all other lines (Fig. 3H). *Hexb* mRNA was strongly reduced in *Hexb^CreER/CreER^* microglia (Fig. 3I), which is consistent with the original report on the generation of *Hexb^CreER^* mice (Masuda et al., 2020). To note, the same study showed no effect on *Hexb* mRNA levels in heterozygous *Hexb^CreER/+^* mice (Masuda et al., 2020). We further assessed *CreER* expression across the lines. Interestingly, despite differences in recombination efficiency, *CreER* expression was similar in *Cx3cr1^YFP-CreER/+ (Litt)^, Cx3cr1^CreER/+ (Jung)^*, and *Tmem119^CreER/+^* microglia, while *CreER* expression was significantly lower in *Hexb^CreER/CreER^* microglia (Fig. 3J). These data suggest that there are some subtle differences in gene expression between Cre lines that should be taken into consideration and that the level of *CreER* RNA is not the best predictor of recombination efficiency across the various CreER lines. It remains to be determined if these differences in *CreER*, *Hexb*, or *Tmem119* RNA are reflected at the level of protein. Indeed, mice homozygous null for *Hexb* develop a lethal neurodegenerative disease (Sango et al., 1995), which we and others did not observe in the *Hexb^CreER/CreER^* mice with reduced *Hexb* mRNA (Masuda et al., 2020). These data suggest that the protein and its enzymatic activity are still preserved in *Hexb^CreER/CreER^* mice despite significantly reduced amounts of mRNA.

**Figure 3.**
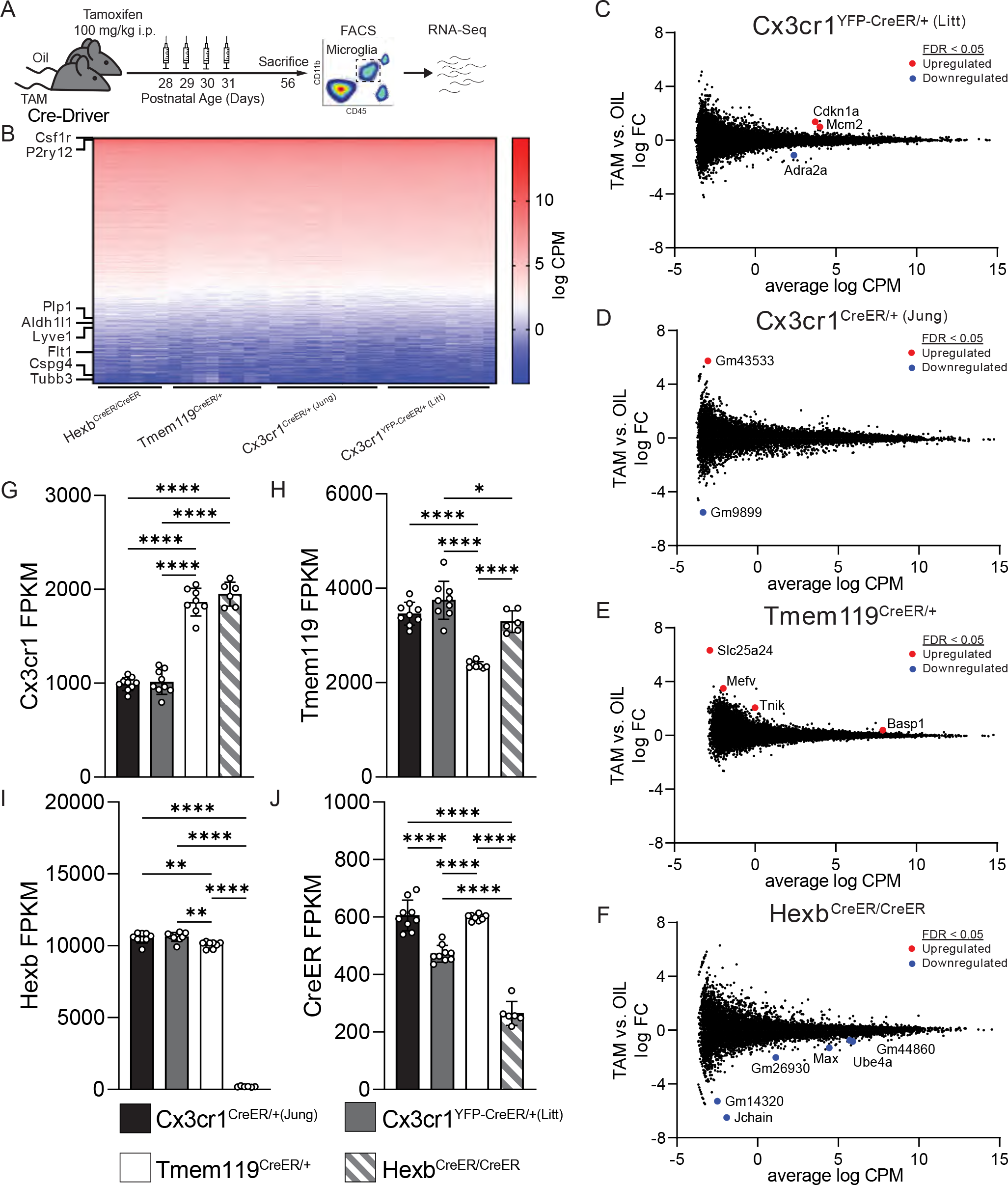
Subtle differences in gene expression exist across microglial CreER lines. **A** Diagram of experimental protocol used to perform RNA sequencing on microglia from CreER lines injected with tamoxifen (TAM) or oil. **B** Heatmap of gene expression values (log counts per million (CPM)) across all samples. Rows corresponding to cell-type specific genes markers for microglia (*Csf1r, P2ry12*), oligodendrocytes (*Plp1*), astrocytes (*Aldh1l1*), border-associated macrophages (*Lyve1*), endothelial cells (*Flt1*), oligodendrocyte precursor cells (*Cspg4*), and neurons (*Tubb3*) are annotated. **C-F** Smear plots of TAM vs. oil injected *Cx3cr1^YFP-CreER/+ (Litt)^* (C), *Cx3cr1^CreER/+ (Jung)^* (D), *Tmem119^CreER/+^* (E), and *Hexb^CreER/CreER^* (F) mice depicting log fold change (FC) on the y-axis against log CPM on the x-axis. Differentially expressed genes with false discovery rate (FDR) < 0.05 are annotated in red (upregulated by TAM) or blue (downregulated by TAM). **G-I** Quantification of genes whose promoters are used to drive CreER expression show that *Cx3cr1* is reduced in *Cx3cr1^YFP-CreER/+ (Litt)^* and *Cx3cr1^CreER/+ (Jung)^* mice. 1-way ANOVA with Tukey’s post hoc. (G, *Cx3cr1^YFP-CreER/+ (Litt)^* vs. *Tmem119^CreER/+^*, *n =* 9, 8 mice, *P* < 0.0001, *q* = 21.02, *df* = 28; *Cx3cr1^YFP-CreER/+ (Litt)^* vs. *Hexb^CreER/CreER^*, *n =* 9, 6 mice, *P* < 0.0001, *q* = 21.29, *df* = 28; *Cx3cr1^CreER/+ (Jung)^* vs. *Tmem119^CreER/+^*, *n =* 9, 8 mice, *P* < 0.0001, *q* = 20.54, *df* = 28; *Cx3cr1^CreER/+ (Jung)^* vs. *Hexb^CreER/CreER^; n =* 9, 6 mice, *P* < 0.0001, *q* = 20.85, *df* = 28) (H, *Tmem119^CreER/+^* vs. *Cx3cr1^YFP-CreER/+ (Litt)^*, *n =* 8, 9 mice, *P* < 0.0001, *q* = 11.64, *df* = 28; *Tmem119^CreER/+^* vs. *Cx3cr1^CreER/+ (Jung)^, n =* 8, 9 mice, *P* < 0.0001, *q* = 14.71, *df* = 28; *Tmem119^CreER/+^* vs. *Hexb^CreER/CreER^, n =* 8, 6 mice, *P* < 0.0001, *q* = 8.876, *df* = 28; *Cx3cr1^CreER/+ (Jung)^* vs. *Hexb^CreER/CreER^*, *n =* 9, 6 mice, *P* < 0.0186, *q* = 4.468, *df* = 28) and (I, *Tmem119^CreER/+^* vs. *Cx3cr1^YFP-CreER/+ (Litt)^*, *n =* 8, 9 mice, *P* < 0.0067, *q* = 5.053, *df* = 28; *Tmem119^CreER/+^* vs. *Cx3cr1^CreER/+ (Jung)^, n =* 8, 9 mice, *P* < 0.0019, *q* = 5.741, *df* = 28; *Tmem119^CreER/+^* vs. *Hexb^CreER/CreER^, n =* 8, 6 mice, *P* < 0.0001, *q* = 94.04, *df* = 28; *Cx3cr1^CreER/+ (Jung)^* vs. *Hexb^CreER/CreER^*, *n =* 9, 6 mice, *P* < 0.0001, *q* = 101.7, *df* = 28; *Cx3cr1^YFP-CreER/+ (Litt)^* vs. *Hexb^CreER/CreER^*, *n =* 9, 6 mice, *P* < 0.0001, *q* = 101.0, *df* = 28). **J** Quantification of *CreER* expression shows significant differences between CreER lines. 1-way ANOVA with Tukey’s post hoc. (*Cx3cr1^CreER/+ (Jung)^* vs. *Cx3cr1^YFP-CreER/+ (Litt)^*, *n =* 9, 9 mice, *P* < 0.0001, *q* = 10.68, *df* = 28; *Tmem119^CreER/+^* vs. *Cx3cr1^CreER/+ (Jung)^, n =* 8, 9 mice, *P* < 0.0001, *q* = 9.830, *df* = 28; *Tmem119^CreER/+^* vs. *Hexb^CreER/CreER^, n =* 8, 6 mice, *P* < 0.0001, *q* = 23.43, *df* = 28; *Cx3cr1^CreER/+ (Jung)^* vs. *Hexb^CreER/CreER^*, *n =* 9, 6 mice, *P* < 0.0001, *q* = 14.95, *df* = 28; *Cx3cr1^YFP-CreER/+ (Litt)^* vs. *Hexb^CreER/CreER^*, *n =* 9, 6 mice, *P* < 0.0001, *q* = 24.51, *df* = 28). Data in G-J are presented as mean ± SEM. See also Figure S3.

### LoxP distance is a key determinant of recombination efficiency in microglial CreER lines

While several parameters (e.g. chromatin accessibility) should be taken into account when predicting recombination efficiency, as longer inter-LoxP distances are correlated with lower recombination rates compared to shorter inter-LoxP distances (Glaser et al., 2005; Van Hove et al., 2020). Since *Rosa26^mTmG^* has a relatively long inter-LoxP distance (2.4 kb), we also wanted to assess how well microglial CreER lines would recombine a relatively shorter inter-LoxP distance. For these experiments, we compared recombination of the *C1qa ^Flox^* allele (1.2 kb inter-LoxP distance) versus the *Rosa26^mTmG^* allele (2.4 kb inter-LoxP distance) using either a highly efficient microglial CreER line *Cx3cr1^YFP-CreER (Litt)^* or a less efficient microglial CreER line *Tmem119^CreER^* line. Unlike initial assessments of recombination across CreER lines in which the *Rosa26^mTmG^* allele was heterozygous, each floxed allele was bred to homozygosity for this comparison. Then, mice were injected with 100mg/kg tamoxifen i.p. or corn oil from P28-P31.

Microglia were collected by FACS at P56, microglial genomic DNA (gDNA) was isolated, and recombination was measured by end-point PCR (Fig. 4A). As predicted, *Tmem119^CreER^* induced more efficient recombination of the floxed allele with a shorter inter-LoxP distance (*C1qa^Flox^*) (Fig. 4B-C) compared to *Rosa26^mTmG^*, which has a longer inter-LoxP distance (Fig. 4D-E).

**Figure 4.**
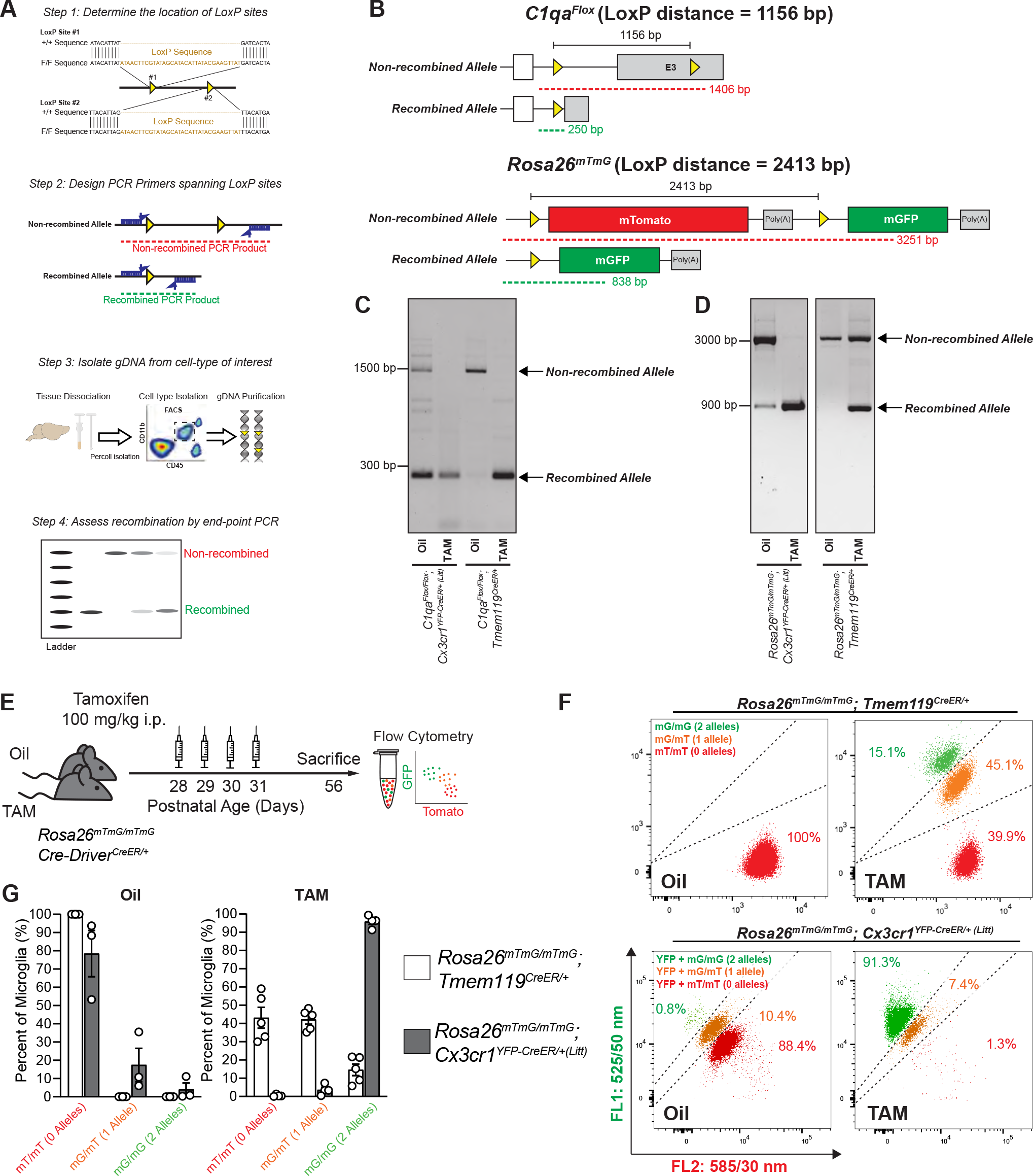
LoxP distance is a determinant of recombination efficiency in microglial CreER lines. **A** Diagram of protocol to assess Cre/LoxP recombination in microglia by end-point PCR of genomic DNA (gDNA). **B** Diagrams of *C1qa^Flox^* allele and the *Rosa26^mTmG^* allele before and after Cre/LoxP recombination show the locations of the LoxP sites (yellow triangles), the LoxP distance, and the end-point PCR products for non-recombined (red) and recombined (green) gDNA. **C-D**End-point PCR of microglial gDNA isolated by florescence-activated cell sorting (FACS) from oil and tamoxifen (TAM) injected *Cx3cr1^YFP-CreER/+ (Litt)^* and *Tmem119^CreER/+^* mice homozygous for (C) *C1qa^Flox/Flox^* or (D) *Rosa26^mTmG/mTmG^*. Gel images of TAM-injected samples from *Cx3cr1^YFP-CreER/+ (Litt)^* mice only show the recombined product for both *C1qa^Flox^* and *Rosa26^mTmG^*, indicating efficient recombination of both alleles. Gel images of TAM-injected samples from *Tmem119^CreER^* mice show only the recombined product for *C1qa^Flox^*, but both recombined and non-recombined products for *Rosa26^mTmG^*, indicating that *Tmem119^CreER^* more efficiently recombines *C1qa^Flox^,* the allele with the shorter LoxP distance. Oil-injected *Cx3cr1^YFP- CreER/+ (Litt)^* mice, but not oil-injected *Tmem119^CreER^* mice, show both non-recombined and recombined products for *C1qa^Flox^* and *Rosa26^mTmG^*, indicating spontaneous recombination. The relative band intensities indicate that *C1qa^Flox^*, the allele with the shorter LoxP distance, undergoes more spontaneous recombination than *Rosa26^mTmG^* in *Cx3cr1^YFP-CreER/+ (Litt)^* mice. **E** Diagram of protocol to assess Cre/LoxP recombination of homozygous *Rosa26^mTmG/mTmG^* microglia isolated by flow cytometry from mice injected with oil or TAM. **F** Flow cytometry analysis of doubly recombined mGFP^+^/mGFP^+^ (mG/mG) vs. singly recombined mGFP^+^/mTomato^+^ (mG/mT) microglia vs. non-recombined mTomato^+^/mTomato^+^ (mT/mT) microglia in *Rosa26^mTmG/mTmG^; Tmem119^CreER/+^* mice with no YFP and *Rosa26^mTmG/mTmG^; Cx3cr1^YFP-CreER/+ (Litt)^* mice expressing YFP. **G** Bar graph of the number of non-recombined mT/mT, singly recombined mG/mT, and doubly recombined mG/mG microglia per animal after injection with oil or TAM. Datapoints represent individual mice. Data is represented as mean ± SEM. See also Figures S4 and S5.

These findings are consistent with previously published work showing nearly 100% recombination at the protein level in *Tmem119^CreER/+^*; *C1qa^Flox/Flox^* mice (Absinta et al., 2021). We also observed efficient recombination of another floxed allele (*Becn1^Flox^*) with an even shorter inter-LoxP distance (0.6 kb) when using the *Tmem119*^CreER^ line (Fig. S4). The *Cx3cr1^YFP-CreER/+ (Litt)^* line efficiently recombined both floxed lines (Fig. 4C, E). End-point PCR was also sufficient to identify spontaneous recombination events in the oil-treated controls of the *Cx3cr1^YFP-CreER (Litt)^* line (Fig. 4C, E) as previously observed by FACS and immunofluorescence microscopy (Figs. 1- 2). These data are also consistent with previous reports of TAM-independent loss of C1QA protein in oil-injected *Cx3cr1^YFP-CreER/+ (Litt)^*; *C1qa^Flox/Flox^* mice (Fonseca et al., 2017). Together, these data suggest that inter-LoxP distance can influence the recombination efficiency of a given CreER line. The *Cx3cr1^YFP-CreER (Litt)^* line has very high efficiency, even at longer inter-LoxP distances, at the expense of spontaneous recombination. In contrast, the *Tmem119^CreER^* line is significantly less efficient for a longer inter-LoxP distances, but this line has a high likelihood of achieving specific and efficient recombination if inter-LoxP distances are shorter than ∼2 kb.

### Detection of multiple independent recombination events in individual microglia

In the process of isolating microglia from *Cx3cr1^YFP-CreER/+ (Litt)^* and *Tmem119^CreER/+^* mice homozygous for the *Rosa26^mTmG^* allele by FACS for end-point PCR (Fig. 4D-E), we noticed 3 populations of microglia for each line (Fig. 4F-H): 1) non-recombined microglia (mTomato^+^/mTomato^+^), 2) microglia with a single allele recombined (mGFP^+^/mTomato^+^), and 3) microglia with two recombined alleles (mGFP^+^/mGFP^+^). These results demonstrate that recombination of the two *Rosa26^mTmG^* alleles in a homozygous *Rosa26^mTmG^*^/^*^mTmG^* microglia occurs independently. This is consistent with previous reports of independent recombination events in individual cells (Cox et al., 2012; Khawaja et al., 2021). In contrast, end-point PCR using gDNA pooled from multiple cells (Fig 4d-e), cannot distinguish the percentage of *Rosa26^mTmG^*^/^*^mTmG^* microglia that are singly recombined vs. doubly recombined. However, we wanted to determine whether we could use endpoint PCR to predict the number of recombined cells detected via the more sensitive FACS assay.

Given that every homozygous cell has 2 alleles of the floxed gene, there is a defined range of uncertainty when estimating the number of fully recombined cells from end-point PCR of gDNA. Using this assumption of 2 alleles per cell, we calculated the theoretical maximum and minimum number of fully recombined cells that would be possible from the recombination of a given number of alleles (Fig. S5A-B). As a test of these assumptions, we compared the percentage of doubly recombined *Rosa26^mTmG^*^/^*^mTmG^* microglia (mGFP^+^/mGFP^+^) detected in each sample by flow cytometry versus the percentage of recombined *Rosa26^mTmG^* alleles (mGFP^+^) inferred from the relative numbers of singly and doubly recombined microglia. This percentage of recombined *Rosa26^mTmG^* alleles (100% for a mGFP^+^/mGFP^+^ cell, 50% for a mGFP^+^/mTomato^+^ cell) is representative of what would be measured by end-point PCR of pooled gDNA. For *Rosa26^mTmG/mTmG^* mice recombined using microglial Cre lines, we found that the number of doubly recombined cells fit along a curve proportional to the square of the *Rosa26^mTmG^* allelic recombination rate (Fig. S5B; *r*^2^ = 0.999). The alignment of observed data with the predicted squared relationship (y=x^2) suggests that the two *Rosa26mTmG* alleles within a given microglia recombine independently, consistent with previous reports (Cox et al., 2012; Khawaja et al., 2021). These data suggest the end-point PCR recombination rate can be used to predict the number of doubly recombined cells for the *Rosa26mTmG* line. However, we cannot assume that all floxed alleles follow the same pattern as *Rosa26mTmG*, and only the theoretical max and min can be used as an appropriate range of uncertainty to use when estimating the percentage of fully recombined cells using gDNA pooled from multiple cells without further experimental confirmation (i.e. with the Rosa26 reporter). Importantly, the theoretical minimum of fully recombined cells nears 100% at high allelic recombination rates, while the theoretical maximum nears 0% with no allelic recombination. Thus, gDNA endpoint PCR can provide a reliable confirmation of complete recombination or complete lack of recombination in homozygous floxed mice. As we and others have shown nearly 100% recombination at the protein level for alleles with shorter inter-LoxP distances with the Tmem line or for most floxed alleles using the efficient Cx3cr1 lines (Absinta et al., 2021; Fonseca et al., 2017), endpoint PCR can be used in these settings effectively. Endpoint PCR can also distinguish spontaneous recombination events in oil-treated controls. However, it is important to note that if a floxed gene is essential for cell survival, this estimation will not accurately reflect the degree of recombination as homozygous recombined cells will die prior to analysis.

### A qPCR protocol to quantitatively assess recombination in microglia

For alleles which do not fully recombine, a more accurate quantification of the number of recombined alleles is needed to estimate the number of fully recombined cells. This is not possible with end-point PCR. Southern blot is a classic and highly quantitative technique, but it requires large amounts (∼5-10 µg) of gDNA(Southern, 2006). We, therefore, implemented a more quantitative measurement of recombination using qPCR (Fig. 5A). For a given floxed mouse line, LoxP sites are first mapped and then three qPCR primer pairs are designed: one outside the LoxP region as a control, a second overlapping a single LoxP region to measure unrecombined DNA, and a third spanning the LoxP region to measure recombined DNA. qPCR conditions are then optimized for the primers, microglial are isolated by FACS, and gDNA is purified for qPCR. We used *Cx3cr1^YFP-CreER (Litt)^*; *Rosa26^mTmG/+^* mice as proof-of-concept to establish this protocol as we also had a well-established FACS protocol for this line. First, we tested the protocol in cultured microglia. As cultured microglia do not efficiently recombine with TAM or 4-OHT, we induced recombination in primary mixed glia cultures, which contain a mixture of microglia and other glial cell types. To induce different degrees of recombination, primary mixed glia prepared from *Cx3cr1^YFP-CreER (Litt)^*; *Rosa26^mTmG/+^* mice were treated with increasing concentrations of 4-OHT or vehicle (ethanol) (Fig. 5B). Microglia were then isolated from the mixed glia and re-plated. Endogenous fluorescence within the purified microglia cultures confirmed recombination by an increase in mGFP and loss of mTomato in 4-OHT-treated cultures (Fig. 5C). Subsequently, cultures were analyzed by flow cytometry to quantify the percentages of microglia that were mGFP^+^ (recombined) or mTomato^+^ (unrecombined) (Fig. 5D). Note, because the mice used for this study were heterozygous for *Rosa26^mTmG/+^*, no singly recombined cells were observed. As expected, 4-OHT-treated cultures had a higher recombination rate in microglia than vehicle-treated cells (Fig. 5E) and recombination efficiency calculated by qPCR was highly correlated with the recombination efficiency calculated by flow cytometry (Fig. 5F; *r*^2^ = 0.954).

**Figure 5.**
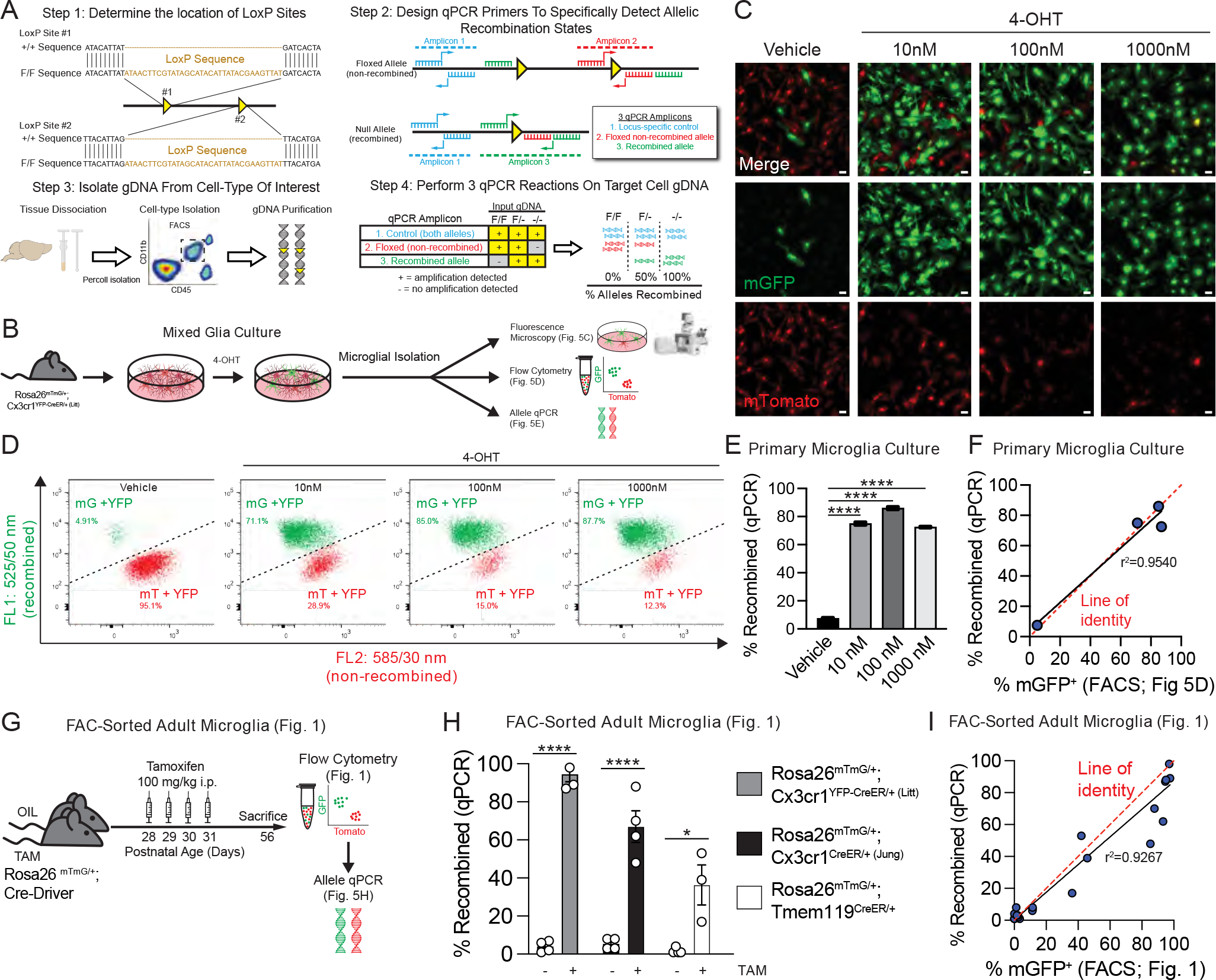
A qPCR protocol to quantitatively assess recombination in microglia. **A** Diagram of protocol used to quantify Cre/LoxP recombination of microglial genomic DNA (gDNA) by quantitative PCR (qPCR). **B** Diagram of experiment to assess Cre/LoxP recombination in primary microglia cultures from *Rosa26^mTmG/+^; Cx3cr1^YFP-CreER/+ (Litt)^* mice after exposure to 4-hydroxytamoxifen (4-OHT). **C** Fluorescent images of endogenous mGFP (green) and endogenous mTomato (red) in primary microglia cultures from *Rosa26^mTmG/+^; Cx3cr1^YFP- CreER/+ (Litt)^* mice after exposure to 4-OHT. Scale bars=25 µm. **D** Flow cytometry analysis of recombined mGFP^+^ (mG) vs. non-recombined mTomato^+^ (mT) microglia in primary microglia cultures from *Rosa26^mTmG/+^; Cx3cr1^YFP-CreER/+ (Litt)^* mice after exposure to 4-OHT. **E** Quantification of the percentage of recombined gDNA by qPCR shows increased recombination in primary microglia exposured to 10 nM, 100 nM, or 1000 nM of 4-OHT (1-way ANOVA with Dunnett’s post-hoc; vehicle vs. 10 nM, *n* = 2 independent cultures, *P* < 0.0001, *q* = 60.37, *df* = 4; vehicle vs. 100 nM, *n* = 2 independent cultures, *P* < 0.0001, *q* = 70.21, *df* = 4; vehicle vs. 1000 nM, *n* = 2 independent cultures, *P* < 0.0001, *q* = 58.14, *df* = 4). **F** Graph of percent recombination of *Rosa26^mTmG^* in primary microglia from *Rosa26^mTmG/mTmG^; Cx3cr1^YFP-CreER/+ (Litt)^* mice after exposure to 4OH tamoxifen as measured by flow cytometry analysis vs. the recombination rate as measured by qPCR of microglial gDNA isolated by fluorescence-activated cell sorting (FACS). Data points fit to a linear curve (black line; r^2^ = 0.9540), closely aligned with the line of identity (red dashed line), indicating that qPCR provides a linear, quantitative measurement of *Rosa26^mTmG^* recombination in *in vitro* samples. **G** Diagram of experiment to assess Cre/LoxP recombination in mice injected with tamoxifen (TAM) or oil. **H** Quantification of the percentage of recombined gDNA by qPCR shows increased recombination in TAM vs. oil for all three CreER lines (*Rosa26^mTmG/+^; Cx3cr1^YFP-CreER/+ (Litt)^*: Student’s t-test, *n* = 4 mice, *P* < 0.0001, *t* = 21.38, *d f*= 6; *Rosa26^mTmG/+^; Cx3cr1^CreER/+ (Jung)^*: Student’s t-test, *n* = 4 mice, *P* = 0.0003, *t* = 7.26, *df* = 6; *Rosa26^mTmG/+^; Tmem119^CreER/+^*: Student’s t-test, *n* = 4 oil, 3 TAM mice, *P* = 0.0111, *t* = 3.925, *df* = 5). **I** Graph of percent recombination of *Rosa26^mTmG^* in microglia in mice injected with TAM or oil as measured by flow cytometry analysis (see also Fig. 1) vs. the recombination rate as measured by qPCR of microglial gDNA isolated by FACS. Data points fit to a linear curve (black line; r^2^ = 0.9267), closely aligned with the line of identity (red dashed line), indicating that qPCR provides a linear, quantitative measurement of *Rosa26^mTmG^* recombination in *in vivo* samples. Data in e and h are presented as mean ± SEM.

After initial validation *in vitro*, we next tested the qPCR protocol on microglia acutely isolated by FACS from *Rosa26*^mTmG/+^; *Cx3cr1^YFP-CreER/+ (Litt)^*, *Rosa26*^mTmG/+^; *Cx3cr1^CreER/+ (Jung)^*, or *Rosa26*^mTmG/+^; *Tmem119^CreER/+^* mice (Fig. 5G). Similar to *in vitro* experiments, the measurements of Cre/LoxP recombination by qPCR from gDNA strongly correlated with recombination efficiency calculated by flow cytometry for all three microglial CreER lines (Fig. 5H-I, Fig. 1E, H; *r*^2^ = 0.9267). Note, the experiments described above were conducted in mice heterozygous for the floxed *Rosa26*^mTmG^ allele. In mice homozygous for the floxed allele, the calculation described in Fig. S5 is needed to estimate the number of fully recombined cells from the recombination rate of gDNA. This calculation is particularly important in experiments when full recombination is needed, such as when deleting a gene in a conditional knockout mouse.

Together, these results provide important proof-of-concept that using gDNA isolated from microglia *in vitro* or *in vivo*, combined with qPCR to detect the floxed allele cassette, is a highly reliable, quantitative method for measuring recombination efficiency and estimating the percentage of fully recombined cells. This is particularly useful for floxed alleles with larger inter-LoxP distances and/or when antibodies are not available to test recombination on a single cell level by FACS or immunofluorescence microscopy.

## Discussion

We have performed an in-depth comparison of the most widely available CreER lines that target microglia (Table 1). The *Cx3cr1^CreER^* lines more efficiently recombine than *Tmem119^CreER^* and *Hexb^CreER^* lines. However, recombination efficiency was increased when *Tmem119^CreER/+^* mice were crossed to floxed mice with inter-loxP distances <2kb. Also, recombination efficiency can be increased by using homozygous *Tmem119^CreER/CreER^* or *Hexb^CreER/CreER^* mice. Other advantages of using the *Tmem119^CreER^* and *Hexb^CreER^* lines over the *Cx3cr1^CreER^* lines include increased specificity for microglia vs. other long-lived macrophages, which was shown previously by other studies (Goldmann et al., 2016; Kaiser and Feng, 2019; Masuda et al., 2020). Also, we have shown that these lines have decreased rates of spontaneous recombination compared to the *Cx3cr1^CreER^* lines. To further aid in assessing recombination of different floxed alleles which are not easily measured by flow cytometry or immunofluorescence microscopy, we describe a protocol to measure recombination of gDNA purified from microglia by end-point PCR and more quantitatively by qPCR. Based on our findings, we outline a number of key considerations when using Cre/LoxP technology in microglia.

**Table 1.** Comparison of microglia CreER lines

| **Cre-LoxP considerations** | ***Cx3cr1*<sup>YFP-CreER</sup>**<br>**(Litt)** | ***Cx3cr1*<sup>CreER</sup>**<br>**(Jung)** | ***Tmem119*<sup>CreER</sup>** | ***Hexb*<sup>CreER</sup>** |
| --- | --- | --- | --- | --- |
| Microglia specificity of CreER expression | ++<sup>1</sup> | ++<sup>2,3</sup> | ++++<sup>4</sup> | ++++<sup>5</sup> |
| Strength of CreER activity | ++++ | ++++ | ++ | ++ |
| Tamoxifen-independent recombination | + | + | - | - |
| Retained expression of gene driving CreER | ++ | ++ | +++ | + |
| Off-target effects of CreER activation + TAM | +++<sup>6</sup> neonate TAM<br>- P28-P31 TAM | ? neonate not tested<br>- P28-P31 TAM | - neonate TAM<br>- P28-P31 TAM | ? neonate not tested<br>- P28-P31 TAM |
| Effective for long LoxP distances (~2.4 kb) | ++++ | ++++ | ++ | ++ |
| Effective for short LoxP distances (~1.2 kb) | ++++ | ++++ | ++++ | ++++ |

Table comparing microglia CreER lines. Each consideration is rated on a scale from low (-) to high (++++), not tested (?). <sup>1</sup>(Parkhurst et al., 2013), <sup>2</sup>(Yona et al., 2013), <sup>3</sup>(Goldmann et al., 2016), <sup>4</sup>(Kaiser and Feng, 2019), <sup>5</sup>(Masuda et al., 2020), <sup>6</sup>(Sahasrabuddhe and Ghosh, 2022)

### Selecting CreER lines for assessment of microglia

Care should be taken to select the appropriate CreER line for each study. *Cx3cr1^CreER^* lines appear to have the strongest Cre activity compared to *Tmem119^CreER^* and *Hexb^CreER^* lines and allow for the most efficient and homogenous targeting of all microglia. On the other hand, the *Tmem119^CreER^* and *Hexb^CreER^* lines offer distinct advantages. *Tmem119^CreER^* or *Hexb^CreER^* more selectively target microglia vs. border-associated macrophages in the CNS and other longer-lived macrophages throughout the body (Goldmann et al., 2016; Kaiser and Feng, 2019; Masuda et al., 2022; Masuda et al., 2020). *Tmem119^CreER^* or *Hexb^CreER^* will further allow to perform potential in vivo microglia competition experiment between microglia with wild-type versus the deleted gene. Also, if studies require neonatal injection of TAM, the *Cx3cr1^YFP-CreER (Litt)^* mice should be avoided as this line and TAM protocol induces an interferon response in microglia (Sahasrabuddhe and Ghosh, 2022) and downregulation of P2RY12 into adulthood (Figure S3). These results suggest long-lasting alterations in microglia. Although experiments were performed with a different reporter line, it was shown that neonatal TAM injection in *Hexb^CreER^* mice induced recombination up to 85% (Masuda et al., 2020), which offers a viable alternative strategy. Another potential option is to use the new and highly specific microglial split-Cre line if temporal regulation of gene recombination is not required (Kim et al., 2021).

In studies in older mice, or in studies comparing mice with TAM-induced recombination at different ages, *Tmem119^CreER^* and *Hexb^CreER^* may also be preferred due to the high rate of spontaneous recombination in *Cx3cr1^CreER^* mice, which we found progressively increases with age (Fig. 2)(Fonseca et al., 2017). Also, for *Tmem119^CreER^* or *Hexb^CreER^* lines, the inter-LoxP distance can be one factor that can be used to predict recombination efficiency. For at least the *Tmem119^CreER^* line, we found that two floxed alleles with shorter inter-LoxP distances have more efficient recombination. It should be noted, however, that other factors can also affect recombination efficiency, such as chromatin accessibility. Still, we recommend determining the LoxP distance of the floxed allele (if not publicly listed) when designing a study prior to selecting the CreER line to use.

### Selecting appropriate controls for Cre-Lox experiments

When performing Cre/Lox experiments, the following should be considered: 1) the impact of TAM exposure, 2) spontaneous Cre/Lox recombination, 3) the endogenous expression of the gene driving CreER, and 4) off-target effects of CreER activation. To assess spontaneous recombination, oil-injected *Cre-Driver^CreER/+^*; *Floxed Allele^Flox/Flox^* mice are recommended. If spontaneous recombination is low, one should ideally use TAM-injected *Cre- Driver^CreER/+^*; *Floxed Allele^+/+^* mice as controls for TAM-injected *Cre-Driver^CreER/+^*; *Floxed Allele^Flox/Flox^* mice. We note that TAM had minimal impact on microglial gene expression 4 weeks after exposure in adolescent mice; however, it is best practice to control for organism-wide effects of TAM (Donocoff et al., 2020; Zhang et al., 2021). If spontaneous recombination is high, oil injected *Cre-Driver^CreER/+^*; *Floxed Allele^Flox/Flox^* mice should be included to control for the effects of spontaneous recombination on important phenotypes. However, oil injected *Cre- Driver^CreER/+^*; *Floxed Allele^Flox/Flox^* mice controls don’t control for off-target effects of TAM or off-target effects of CreER activation, so TAM injected *Cre-Driver^CreER/+^*; *Floxed Allele^+/+^* mice are also recommended as controls. A high rate of spontaneous recombination is also concerning if parents express both the Cre-Driver and the floxed allele. For example, ablation of a gene in microglia in the parents could affect parental behavior and/or pregnancy. Oil-injected *Cre- Driver^CreER/+^*; *Floxed Allele^Flox/Flox^* and/or TAM-injected *Cre-Driver^CreER/+^*; *Floxed Allele^+/+^* mice littermate controls could be used to control for any effects of spontaneous CreER activation in the parents. Also, if endogenous expression of genes driving the CreER are reduced, as we observed in the CreER lines in this study, a comparison of mice negative for the CreER and positive for *Floxed Allele^Flox/Flox^* should be considered.

### Protocols for assessing recombination in microglia

Lastly, we provide protocols to assess recombination efficiency in microglia by flow cytometry, immunofluorescence microscopy, end-point PCR of gDNA, and qPCR of gDNA. Since Cre/LoxP recombination is dependent on the LoxP distance, using a floxed reporter line as a proxy can be highly misleading. Methods that measure bulk levels of RNA or protein in microglia (e.g. RNA qPCR or Western blot) are useful, but they are best for molecules expressed at higher levels and do not account for independent recombination events. Direct assessment of recombined floxed alleles of interest is best practice. If reliable antibodies exist or expression of a genetic reporter is the desired recombination outcome, recombination can be reliably assessed at the protein level in microglia by flow cytometry or immunofluorescence microscopy in tissue. These approaches are preferred since each cell can be individually classified as recombined or non-recombined at the protein level. Single molecule fluorescence *in situ* hybridization (smFISH) is also an option, although proteins may have a long half-life and these techniques can be costly. Still, an advantage of immunofluorescence microscopy and smFISH is that these techniques allow the assessment of recombination in other cell types in the tissue of interest. However, if the protein or gene cannot be easily detected by these methods, end-point PCR of microglial gDNA is a reliable method, particularly when recombination is near 0 or 100%. qPCR of microglial gDNA offers an even more quantitative measurement and can be predictive of singly and doubly recombined alleles. Southern blotting is likely the most quantitative, but this method requires a large amount of gDNA (Southern, 2006). Still, it should be noted that if a floxed gene is necessary for cell survival, the assessments of recombination will not include dead or dying homozygous cells and the analysis will be strongly biased towards heterozygous cells.

In summary, as interest in microglia continues to grow, common guidelines for the effective use of widely used genetic tools such as Cre/LoxP is critical to achieve reliable, reproducible results. Cre/LoxP technology is of particular importance for molecular studies designed to elucidate the microglia-specific functions of individual genes associated with increased risk of brain diseases such as Alzheimer’s disease and schizophrenia. By providing the pros and cons of different CreER lines, offering guidance on how to properly select the appropriate controls, and implementing best practice protocols for accurately measuring Cre/LoxP recombination in microglia, we aim to assist the field in selecting the best Cre line for their question and in achieving more reliable results with Cre/LoxP recombination in microglia.

Although the specific CreER lines and isolation methods will differ for other cell types, the guidelines and protocols outlined in this study are also informative for the general use of Cre/Lox technology across all other cell types.

## Acknowledgements

This work was supported by NIMH-R01MH113743 (DPS), NIMH- R01 MH118329 (AS), NINDS- R01NS117533 (DPS), NINDS- R01NS106721 (AS), NIA-RF1AG068281(DPS), NIA-RF1AG068558 (AS), NIA- R01AG072489 (AS), the Dr. Miriam and Sheldon G. Adelson Medical Research Foundation (DPS, RK), Autism Speaks Weatherstone Predoctoral Fellowship 11779 (PAF), TM was supported by AMED (JP20gm6310016, JP21wm0425001), JSPS (KAKENHI JP21H02752, JP21H00204, JP22H05062). MP and KPK were supported by the DFG (CRC/TRR167 “NeuroMac”).Schematics were created with Biorender.com.

## Author Contributions

TEF, PAF, AS and DPS designed the study. TEF, PAF, and DPS wrote the manuscript. TEF and PAF performed most experiments and analyzed most data. CO assisted in the design of initial experiments and performed experiments to isolate microglia for assessment of recombination by qPCR and endpoint PCR. RK performed RNA sequencing experiments. HS and AC performed experiments to isolate and extract RNA from microglia from Hexb^CreER^ mice. AS, MP, TM, LA, and KK provided critical input into study design and feedback on writing of the manuscript.

## STAR Methods

### Resource Availability

#### Lead contact

Further information and requests for resources should be directed to and will be fulfilled by the lead contact, Dorothy Schafer.

### Materials availability

This study did not generate new unique reagents.

### Data and code availability

RNA-seq data have been deposited at GEO and are publicly available as of the date of publication. Accession numbers are listed in the key resources table. This paper does not report original code. Any additional information required to reanalyze the data reported in this paper is available from the lead contact upon request.

### Experimental model and subject details

#### Animals

*Cx3cr1^YFP-CreER/+ (Litt)^* (Stock# 021160, B6.129P2(Cg)-Cx3cr1^tm2.1(cre/ERT2)Litt^/WganJ), *Cx3cr1^CreER/+ (Jung)^* (Stock# 020940, B6.129P2(Cg)-Cx3cr1^tm2.1(cre/ERT2)Jung^/J), *Tmem119^CreER/+^* (Stock# 031820, C57BL/6-Tmem119^em1(cre/ERT2)Gfng^/J), Rosa26^mTmG/+^ (Stock# 007676, B6.129(Cg)-Gt(ROSA)^26Sortm4(ACTB-tdTomato,-EGFP)Luo^/J), *C1qa*^Flox^ (Stock# 031261, B6(SJL)-C1qa^tm1c(EUCOMM)Wtsi^/TennJ), and *Becn1^Flox^* (Stock# 028794, Becn1^tm1.1Yue^/J) mice were obtained from Jackson Laboratories (Bar Harbor, ME). *Hexb^CreER/+^* (Hexb^em2(cre/ERT2)Mp^) mice were directly obtained from Dr. Marco Prinz. All mice were maintained on a C57Bl6/J background. Animals were group housed after weaning and maintained on a 12 hour light/dark cycle with food and water provided *ad libitum*. All animals were healthy and were not immune compromised.

Littermates of the same sex were randomly assigned to experimental groups. Equal numbers of males and females were used for all experiments except for RNA Sequencing for which only males were used. Unless otherwise noted, all experiments were performed in 8 week old mice. All animal experiments were performed in accordance with Animal Care and Use Committees (IACUC) and under NIH guidelines for proper animal use and welfare.

### Method Details

#### Tamoxifen injection

To induce CreER activity in adults, animals were injected once daily i.p. with 100 mg/kg tamoxifen Sigma T5648) dissolved 20 mg/mL in corn oil (Sigma C8267) on postnatal days 28, 29, 30, and 31. Neonatal animals were injected once daily with 1 mg/mL tamoxifen on either: days P1-P4 (50 µg) and P14 (100 µg) or days P1, P3, P5, P7 (50 µg).

#### Microglia isolation from adult mouse brain

Microglia were isolated following a published protocol (Hammond et al., 2019) with some modifications. The complete protocol is as follows: All tubes used to spin cells during this protocol were precoated for at least one hour with 1% bovine serum albumin (BSA; Sigma A2153) diluted in ultrapure water to improve cell yields and viability. Centrifuges and tools were all prechilled to 4°C or on ice. When collecting samples for RNA, standard RNAse free conditions were followed. Mice for gDNA sample collection were anesthetized and transcardially perfused using ice cold, Ca^2+^-free, Mg^2+^-free Hank’s balanced salt solution (HBSS; GIBCO) and the brains were quickly dissected and placed in HBSS on ice. Mice used for RNA sample collection were sacrificed with CO2 and the brains were removed and placed in HBSS on ice.

For mice where one hemisphere was used for immunohistochemistry, brains were hemisected along the sagittal midline, the right half was transferred to a 4% paraformaldehyde (PFA; Electron Microscopy Sciences 15711) PBS solution for immunohistochemistry and the left half placed in HBSS for microglial isolation. For microglial isolation, the cerebellum and brainstem were removed and the remaining brain minced using a razor, taking care not to carve plastic fragments from the dish. Samples were then dounce homogenized (2mL homogenizer; DWK Life Sciences (Kimble)) in 1.5mL ice cold HBSS 30 times each with the loose (A) and tight (B) pestles while simultaneously rotating the pestle. The cell suspension was then transferred through a pre-wet (with HBSS) 70-micron cell strainer (Miltenyi 130-098-462) into a prechilled, BSA-coated 50 mL tubes. Cell suspensions were then spun down at 300 xG for 10-minutes in a centrifuge set to 4°C. For microglial enrichment and myelin removal, a 40% Percoll solution was prepared from Percoll (Sigma GE17-0891-01), 10X HBSS, and water for a final 1X HBSS concentration then pH adjusted to 7.3 with dilute HCl. Cell pellets were resuspended in 2mL 40% Percoll then transferred to a pre-chilled, BSA-coated 15mL tube. The 50mL tube was rinsed with up to an additional 7mL 40% Percoll and transferred to the 15mL tube. The final volume for each sample was adjusted to 10mL total by adding Percoll to the 10mL line. The samples were then spun for 30 min at 500 xG with full acceleration and braking. The myelin at the top of the solution was removed using vacuum suction, and then the rest of the Percoll solution. The cell pellet was resuspended in 1-2mL ice-cold HBSS and transferred to a new pre-chilled, BSA-coated 15mL tube. An additional 6mL of HBSS was used to rinse the Percoll spin tube to ensure complete transfer of cells to the new tube. The final volume was adjusted to 10mL with HBSS and spun again for 10 min at 300 xG at 4°C. All samples were then resuspended in 300µL of ice cold FACS buffer (0.5% BSA, 1mM EDTA, in 1x PBS, Sterile Filtered) then transferred to a pre-chilled, BSA-coated 1.5mL tube. Several additional rinses with 300µL per transfer were done to ensure complete transfer of cells from the 15mL tube to the 1.5mL tube. The samples were again spun for 10 min at 300 xG at 4°C and resuspended in ice-cold FACS buffer containing APC anti-CD11b (eBioscience clone M1/70), PerCP-Cy5.5 anti- CD45 (eBioscience clone 30-F11) antibodies at a 1:50 dilution for 15 min on ice. Samples were then washed in 1 mL of ice cold FACS buffer and spun for 10 min at 300 xG at 4°C and then resuspended in 300-500µL of ice cold FACS buffer supplemented with DAPI (Miltenyi 130-111- 570, 1:100 dilution). 10,000-50,000 microglia (DAPI^-^ CD11b^+^ CD45^Mid^) were sorted per sample on a SONY MA800 or SONY MA900 into 1.5mL tubes containing either 200µL FACS buffer for gDNA samples or 300µL Trizol LS reagent (Invitrogen) for RNA samples. For samples from *Rosa26^mTmG^* mice, GFP (488 nm laser; FL1 525/50 nm) and TdTomato (561 nm laser; FL2 585/30 nm) fluorescent signal was also collected for further analysis using FlowJo™ v10.80 Software (BD Life Sciences). The sample and collection tubes were maintained at 4°C and each sample was kept on ice before and after the sort. Each sample took approximately 10-20 min to sort. After the sort, gDNA samples were spun at 10,000 xG for 10 minutes and the supernatant was discarded. Cell-pellet-containing tubes were kept at -20°C for short term storage until all samples were collected for batch processing (see genomic DNA isolation). The RNA samples (Trizol tubes) were stored at -80°C until all samples were collected for batch processing (see total RNA isolation).

#### Immunostaining & confocal microscopy

Brains from transcardially perfused mice were fixed in 4% paraformaldehyde (PFA, electron microscopy sciences) 1X PBS solution. Brains were kept at 4°C overnight then rinsed with PBS before being transferred to 30% sucrose 1X PBS solution for dehydration. 40µm thick sagittal sections were cut on a vibratome (Leica) at in a bath containing PBS. Sections were transferred to individual wells of 24-well plates for staining. The sections were blocked for 1 hour using PBTGS (0.1M phosphate buffer, 5% normal goat serum (NGS; Sigma G9023), 0.3% Tritonx- 100 (Sigma X100)). Sections were incubated overnight at room temperature (RT) with primary antibodies diluted in PBTGS. Primary antibodies used: chicken anti-GFP (1:1000, Abcam ab13970), rabbit anti-P2RY12 (1:1000, anaspec AS-55043A), and rabbit anti-IBA1 (1:500, Wako 019-19741). Sections were washed 3x in 0.1M PB for 5-minutes each. Sections were incubated for 2-3 hours at RT with secondary antibodies diluted in PBTGS. Secondary antibodies used: goat anti-chicken-AlexaFluor488 (1:1000, Life Technologies), goat anti-rabbit- AlexaFluor594 (1:1000, Life Technologies), and anti-rabbit-AlexaFluor647 (1:1000, Life Technologies). Sections were washed 3x in 0.1M PB for 5-minutes each. Next, sections were mounted onto a glass slide then allowed to dry. Fluoroshield mounting media containing DAPI (Sigma F6057) was applied to the section and a coverslip was added. The slide was sealed with nail polish along the edge. Sections were imaged on a Zeiss Observer microscope (Zeiss; Oberkochen, Germany) equipped with 405 nm, 488 nm, 555 nm, and 639 nm lasers and Zen black acquisition software (Zeiss; Oberkochen, Germany).

#### Total RNA Isolation

Total RNA was purified from FACS sorted microglia. Samples were kept on ice unless otherwise stated. Samples were retrieved from the -80°C freezer and allowed to thaw on ice, then transferred a 2mL phaselock tube. We collected ∼50,000 microglia, which had a sort volume of ∼140 µL, into 300 µL Trizol. To reach the ideal Trizol:sample ratio of 3:1, we adjusted the volume of Trizol to 420µL by adding 120µL Trizol to the phaselock tube. The samples were then incubated at room temperature (RT) for 5-minutes. Next, 110µL of chloroform: isoamyl alcohol (Sigma) was added to each tube and then the tube was shaken vigorously. Samples were incubated again at RT for 5-minutes with repeated shaking. Next, samples were spun at 13,000 xG for 10-minutes at 4°C. Next, an additional 110µL of chloroform: isoamyl alcohol was added to the tube and then the tube was shaken vigorously. The samples were then spun again at 13,000 xG for 10-minutes at 4°C. After the spin, 300µL of the aqueous volume from above the gel was transferred to a clean 1.5mL tube. Next, 5µL glycoblue (Invitrogen), and 30µL 3M Sodium Acetate pH5.5 (Ambion) were added to the tube and mixed with gently tapping to ensure fully mixed. Next, 300µL isopropanol (Sigma) was added and the tubes were mixed by inverting 10 times. The samples were transferred to the -80°C for at least one night. The next day, sample were retrieved from the -80°C freezer and spun for 30-minutes at 13,000 xG and 4°C. The supernatant was discarded and 1mL of ice-cold 75% ethanol was added to wash the pellet. With careful pipetting, the pellet was dislodged from the bottom of the tube. Samples were spun again at 13,000 xG at 4°C for 10 minutes then rinsed with ice-cold 75% ethanol for a second time. Next the samples were spun for the final time at 13,000 xG at 4°C for 10 minutes. After the spin, the supernatant was discarded and the tubes were placed at RT with the caps open to air-dry for 10-15 minutes. Dried pellets were resuspended in 20µL nuclease-free ultrapure water. Samples were then stored at -80°C until shipped for quality control and further processing for RNAseq. Samples were prepared in batches containing all tamoxifen and oil injected samples from a particular CreER line.

#### RNA Sequencing

RNA integrity was assessed on an Agilent 2100 Bioanalyzer and samples with RIN < 7 were excluded. Libraries were prepared from the remaining samples using SMART-Seq v4 + Nextera XT and then sequenced on NovaSeq 6000 (Illumina) with PE 100X2. Quality control (QC) was performed on base qualities and nucleotide composition of sequences, to identify problems in library preparation or sequencing. Short reads were aligned using STAR (v2.4.0) to the mouse genome (mm10). Average input read counts were 87.3M per sample and average percentage of uniquely aligned reads was 87.5%. Read counts for mouse refSeq genes were generated by HT-seq (v0.6.1) (Anders et al., 2015). Low count transcripts were filtered, and count data were normalized using the method of trimmed mean of M-values (TMM). Differentially expressed genes (FDR < 0.05) were then identified using the Bioconductor package EdgeR (Robinson et al., 2010).

#### Genomic DNA Isolation

To purify genomic DNA (gDNA) from microglia, cells were lysed with 50µL Bradley Lysis Buffer (10mM Tris-HCl, 10mM EDTA, 0.5% SDS, 10mM NaCl). Samples were incubated at 55°C for 3- hours then returned to room temperature. Next 2µL 5M NaCl (Ambion) was added to each tube and mixed with finger flicking. Next 1µL glycogen (ThermoFisher Scientific) was added to each tube and mixed. Finally, 125µL 100% ethanol was added to each tube and mixed by inverting the tubes 5-10x. The samples were then incubated at -20°C for at least one hour to allow for gDNA precipitation. To pellet the gDNA precipitant, samples were spin at 20,000 xG, 4°C for 30- minutes. The supernatant was removed and the pellet was washed with ice-cold 70% ethanol, vortexed on medium then spin again at 15,000 xG. 4 for 10-minutes. This washing step was repeated two additional times for a total of three washes. After the final spin, the supernatant was removed and the pellet briefly dried using a speed-vac. The pellet was then resuspended in 30µL ultrapure water, pre-warned to 50-55°C. Samples were then incubated at 55°C for at least 30-minutes to facilitate resuspension. The concentration of DNA was determined using a Nanodrop (Thermo Scientific). Samples were stored at -20°C for short term storage and -80°C for long-term storage.

#### LoxP Site Mapping

Benchling was used for all primer design (https://www.benchling.com/). Putative LoxP site locations were obtained by referencing the original published paper or searching the MGI database (http://www.informatics.jax.org/). The benchling primer wizard was then used to design three primer pairs with a high probability of flanking the second loxP site: one primer in the exon, one primer in the intron containing the loxP site. DNA primers were obtained from integrated DNA technology, stored at -20°C 100 µM, and used at 20 µM. We tested the primer pairs by running a PCR using wild type (+/+) and homozygous floxed (F/F) gDNA with Apex 2X Taq RED Master Mix with 1.5mM MgCl2. The PCR products were run out on a 1.5% agarose gel at 100V (constant voltage) for 30-45 minutes. A picture of the gel was taken and the best primer pair per loxP site was identified. Next, we repeated the PCR with successful primer pairs in replicate (3-4) to then pool replicates. Next, we ran pooled reactions (20µL) on a gel using low melting temp agarose. Next, we took pictures of the gel then cut out the +/+ and F/F bands from the gel with a razor under UV light using proper eye protection. Next, we used the Qiagen gel extraction kit (Qiagen) to purify the amplicons following the manufacturers protocol. DNA is eluted from the Qiagen spin column with 30µL Buffer EB (10mM Tris-Cl, pH 8.5). A Nanodrop (Thermo Scientific) was used to determine the DNA concentration. Samples were submitted to GeneWiz for sangar sequencing. Each amplicon was sequenced in both directions using the forward and reverse primers separately. The sequence results were entered into benchling and using the alignment function, the +/+ and F/F sequences were aligned to locate the loxP sequence (ATAACTTCGTATAGCATACATTATACGAAGTTAT). The sequence directly up- and downstream of the loxP sequence was used to map the coordinates of the loxP in the genome. The distance between the loxP sites in base pairs (bp) was determined in benchling.

#### End-point PCR

Primer pairs flanking both LoxP sites were generated used Benchling. Primer pairs were screened by PCR on non-recombined and recombined DNA. For longer PCR products, Hot Start VeriFi Taq polymerase (PCR Biosystems) was used. Gel electrophoresis was performed on 1.5% agarose gels at 100 V to distinguish PCR products of different sizes. Once the best primer pair was selected, PCR was performed on gDNA from FACS microglia samples to estimate the percentage of DNA recombination. Gel images were collected using a Bio-Rad ChemiDoc.

#### Primary microglia cultures

Mixed glia cultures were generated from P1 *Rosa26^mTmG/+^; Cx3cr1^YFP-CreER/+ (Litt)^* mice and grown in advanced DMEM/F12 culture medium supplemented with fetal bovine serum (10%, GIBCO), pen/strep (GIBCO), and glutaMax (GIBCO) at 37°C 5% CO2 for two weeks. Mixed glia cultures were maintained for two weeks with 50% media changes after 5-7 days. CreER activity was induced by addition of 4-hydroxy-tamoxifen (4-OHT, Sigma) dissolved in ethanol to the mixed glia cultures on DIV 13. Three final concentrations were tested: 10nM, 100nM, and 1000nM. Loosely adhered microglia were shaken off the at 300rpm for 3 hours at 37°C, 5% CO2 and plated in 24-well plates at a density of 60,000 cells per well 72-hours after tamoxifen application. The next day, the media was completely removed and changed to serum-free microglia growth media. Microglia growth media consists of neurobasal media (GIBCO), insulin (Sigma-Aldrich), sodium pyruvate (GIBCO), pen/strep (GIBCO), thyroxine (Sigma-Aldrich), glutamax (GIBCO), B27 (GIBCO), n-acetyl cysteine (Sigma-Aldrich), m-CSF (Shenandoah), transferrin (Sigma-Aldrich), bovine serum albumin (Sigma-Aldrich), putrescinne (Sigma-Aldrich), progesterone (Sigma-Aldrich), and sodium selenite (Sigma-Aldrich). After three days in serum-free conditions, the entire media was changed. Fluorescent images of the primary microglia were acquired 24- hours after changing to microglia growth media using an Observer Spinning Disk Confocal microscope equipped with a 40x objective, diode lasers (405 nm, 488 nm, 594 nm, 647 nm) and Zen Blue acquisition software (Zeiss; Oberkochen, Germany). Genomic DNA (gDNA) was also collected 72-hours after tamoxifen induction.

#### Quantitative PCR

Quantitative PCR (qPCR) was performed using primers designed in benchling (https://www.benchling.com/) to generate specific PCR products for 1) a control region, 2) non-recombined gDNA and 3) recombined gDNA. For each primer pair, a 2-fold dilution series was generated with gDNA and only primer pairs with linear amplification were used for qPCR analyses. Primer stocks were prepared at 100µM then further diluted to 20µM working concentration. The final primer concentration in the PCR reaction is 400nM. Each gDNA sample was tested for the three different qPCR primer pairs (control region, non-recombined allele, recombined allele). PowerSYBR master mix (Life technologies) was used and the plates were run using a CFX96 real time PCR instrument (BioRad). A standard curve for each primer pair was run on recombined or non-recombined gDNA and the equation of the best-fit line was used to extrapolate the abundance of the allele in each sample from the cycle number.

### Quantification and Statistical Analysis

Statistical analyses were performed using GraphPad Prism 9.4.1. The exact number of samples (*n*), what *n* represents, the statistical tests used, and *p* values for each experiment are noted in the figure legends. For comparisons between two groups, we used Student’s t-test. For comparisons between more than two groups, we used 1-way ANOVA followed by Tukey’s post-hoc test between groups. For 2-way comparisons, we used 2-way ANOVA followed by Sidak’s post-hoc test.

---

**Figure S1.**
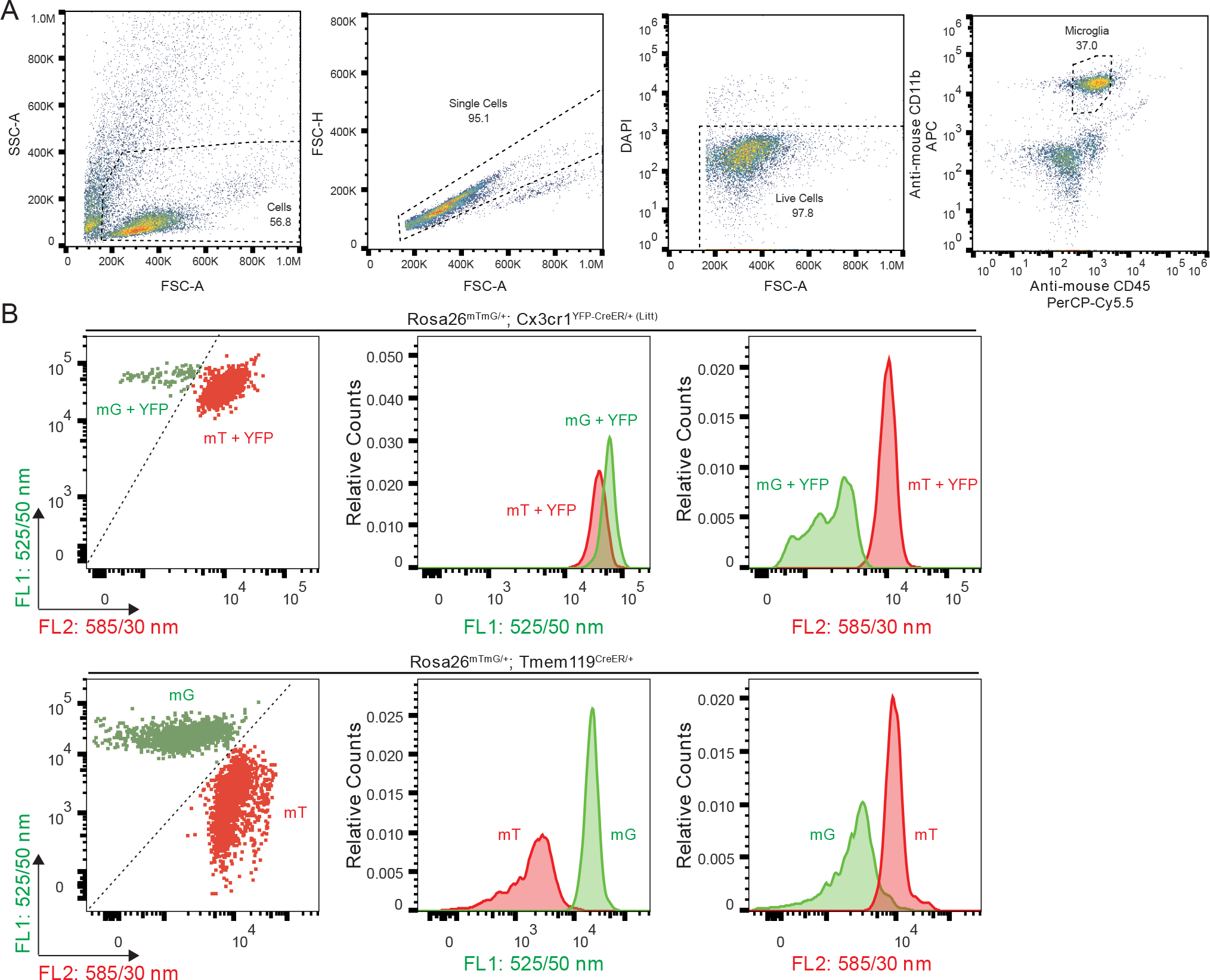
Flow cytometry of *Rosa26^mTmG^* microglia. Related to Figure 1 **A** Plots of flow cytometry gates used for fluorescence-activated cell sorting (FACS) of microglia. **B** Flow cytometry analysis of recombined mGFP^+^ (mG) vs. non-recombined mTomato^+^ (mT) microglia in *Rosa26^mTmG/+^; Cx3cr1^YFP-CreER/+ (Litt)^* expressing YFP and *Rosa26^mTmG/+^; Tmem119^CreER/+^* mice with no YFP. In both CreER lines, the recombined microglia form a distinct population identified by reduced fluorescence in the FL2 585/30 nm channel and increased fluorescence in FL1 525/50 nm channel.

**Figure S2.**
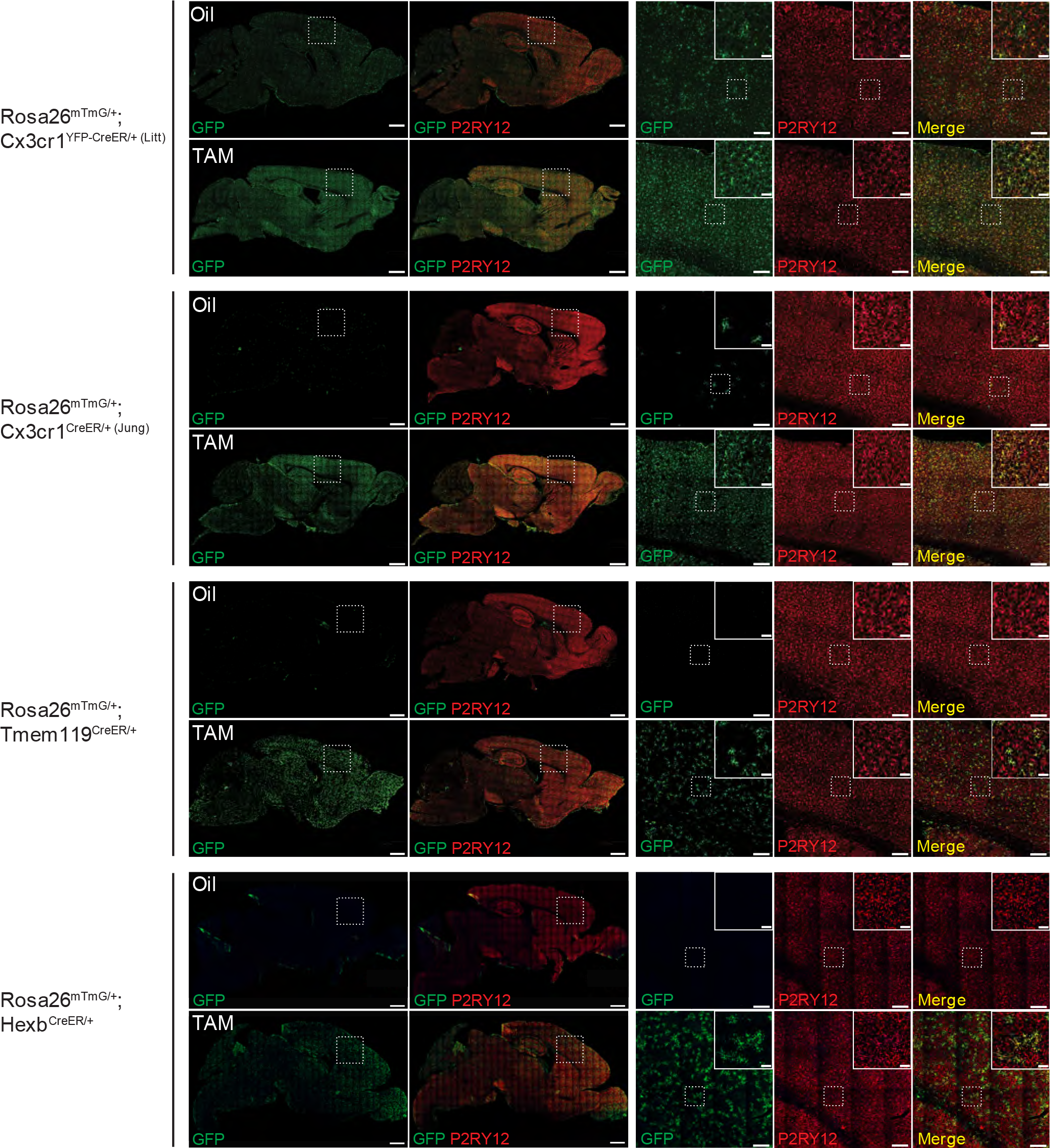
Immunohistochemistry for GFP validates flow cytometry analysis of *Rosa26^mTmG^* recombination. Related to Figure 1 Representative immunofluorescent images of brain sections from right hemisphere of oil and tamoxifen (TAM) injected mice (see also Figs. 1 and 2). Sections were immunolabeled for anti- P2RY12 (red) to identify microglia and anti-GFP (green) to identify recombined cells. The number of recombined mGFP^+^ microglia (white arrows) matches the results observed by flow cytometry (see Fig. 1). In the *Cx3cr1^YFP-CreER/+ (Litt)^* line, the soma of unrecombined microglia are also immunolabeled by anti-GFP due to the constitutive expression YFP. Scale bars 1 mm (full image), 200 µm (large inset), 50 µm (small inset).

**Figure S3.**
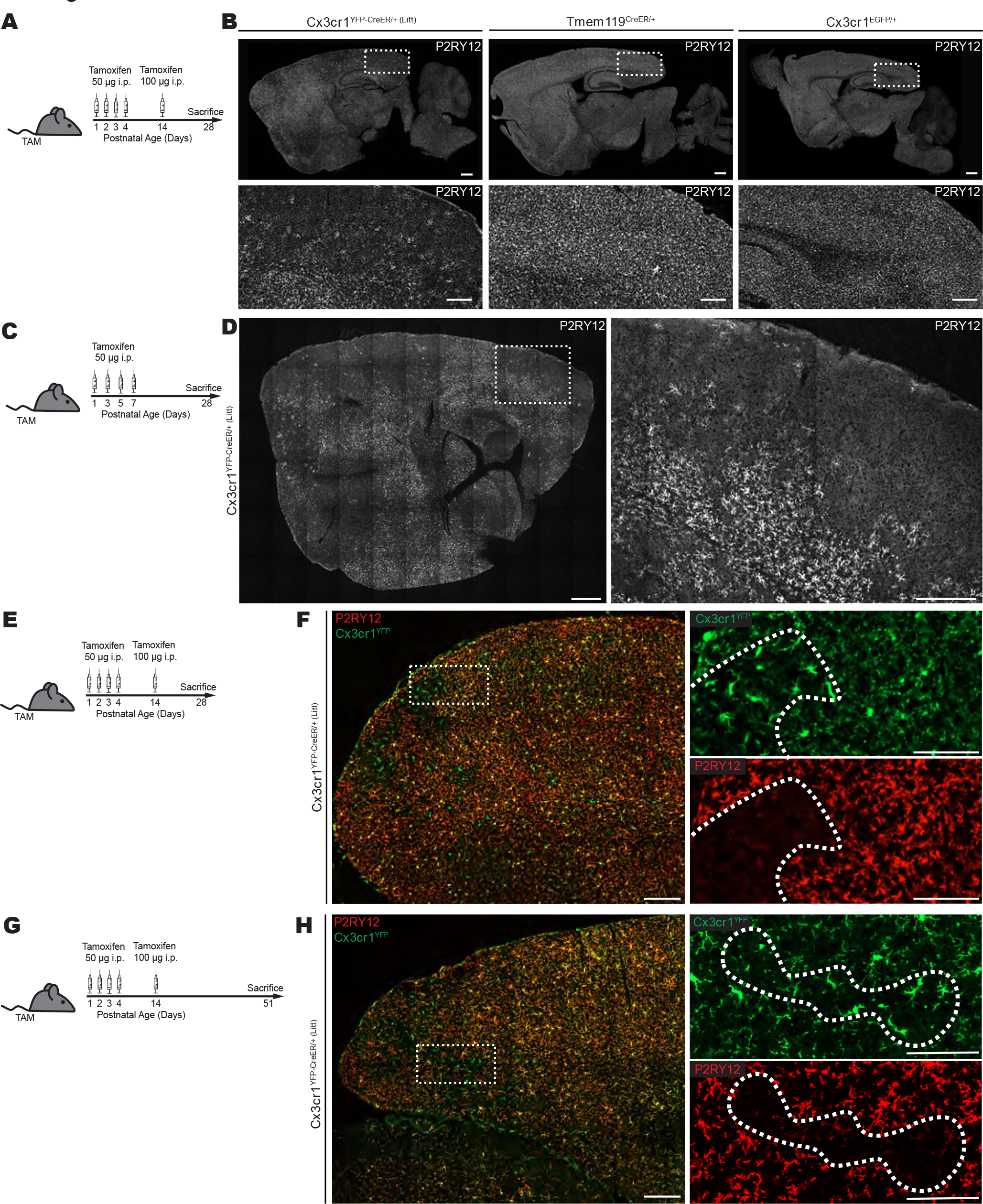
Neonatal CreER activation induces loss of homeostatic microglia in *Cx3cr1^YFP-^ CreER/+ (Litt)* _mice_. Related to Figure 3 **A** Timeline of neonatal tamoxifen (TAM) injection for images in (B). **B** Fluorescent images of brain sections from TAM injected *Cx3cr1^YFP-CreER/+ (Litt)^*, *Tmem119^CreER/+^,* and *Cx3cr1^EGFP/+^* mice, immunolabeled with anti-P2RY12. Large regions of the cortex are devoid of anti-P2RY12 immunofluorescence in *Cx3cr1^YFP-CreER/+ (Litt)^* mice, but not *Tmem119^CreER/+^,* or *Cx3cr1^EGFP/+^* mice. Scale bars 500 µm. Insets 200 µm. **C** Timeline of neonatal TAM injection for images in (D). **D** Fluorescent images of brain sections from TAM injected *Cx3cr1^YFP-CreER/+ (Litt)^* mice, immunolabeled with anti-P2RY12. Large regions of the cortex are devoid of anti-P2RY12 immunofluorescence. Scale bars 500 µm. Insets 200 µm. **E** Timeline of neonatal TAM injection for images in (F). **F** Fluorescent images of brain sections from TAM injected *Cx3cr1^YFP-CreER/+ (Litt)^* mice, immunolabeled with anti-P2RY12 (red) and anti-GFP to identify *Cx3cr1^YFP+^* cells (green). Regions of the cortex devoid of anti-P2RY12 immunofluorescence (dotted outline in inset) contain *Cx3cr1^YFP^*^+^ cells with a morphology characteristic of reactive microglia. Scale bars 500 µm. Insets 100 µm. **G** Timeline of neonatal TAM injection for images in (H). **H** Fluorescent images of brain sections from TAM injected *Cx3cr1^YFP-CreER/+ (Litt)^* mice, immunolabeled with anti- P2RY12 (red) and anti-GFP to identify *Cx3cr1^YFP+^* cells (green). Regions of the cortex devoid of anti-P2RY12 immunofluorescence (dotted outline in inset) contain *Cx3cr1^YFP^*^+^ cells with a morphology characteristic of reactive microglia. Scale bars 500 µm. Insets 100 µm.

**Figure S4.**
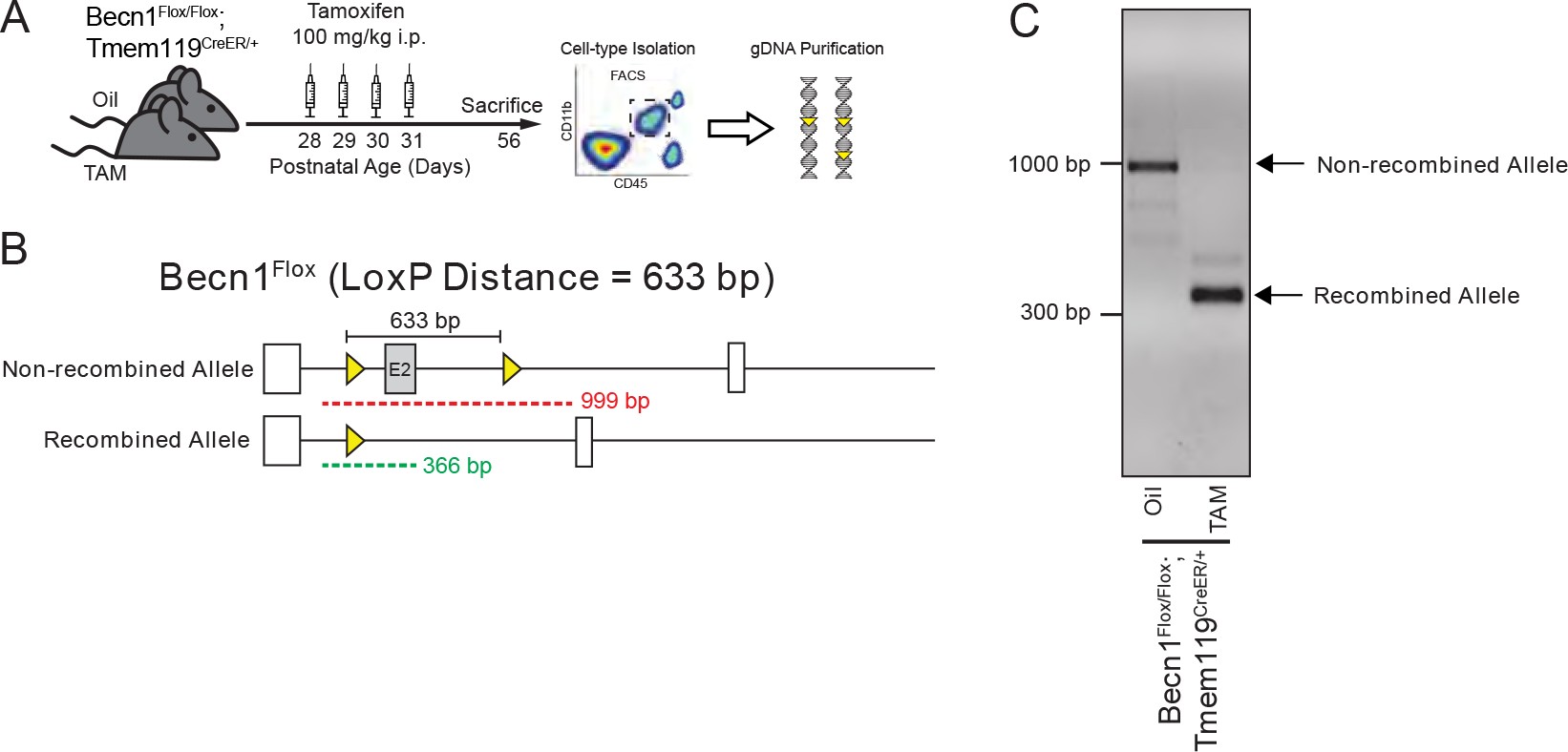
*Tmem119^CreER^* efficiently recombines short LoxP distances. Related to Figure 4 **A** Diagram of protocol to assess Cre/LoxP recombination of genomic DNA (gDNA) from microglia isolated by fluorescence-activated cell sorting (FACS) from *Becn1^Flox/Flox^*; *Tmem119^CreER/+^* mice injected with oil or tamoxifen (TAM). **B** Diagram of *Becn1^Flox^* allele before and after Cre/LoxP recombination showing the locations of the LoxP sites (yellow triangles), the LoxP distance, and the end-point PCR products for non-recombined (red) and recombined (green) gDNA. **C** Gel images of oil-injected samples from *Becn1^Flox/Flox^*; *Tmem119^CreER/+^* mice only show the non-recombined allele and TAM-injected samples from *Becn1^Flox/Flox^*; *Tmem119^CreER/+^* mice only show the recombined allele, indicating no spontaneous recombination without TAM and efficient recombination with TAM.

**Figure S5.**
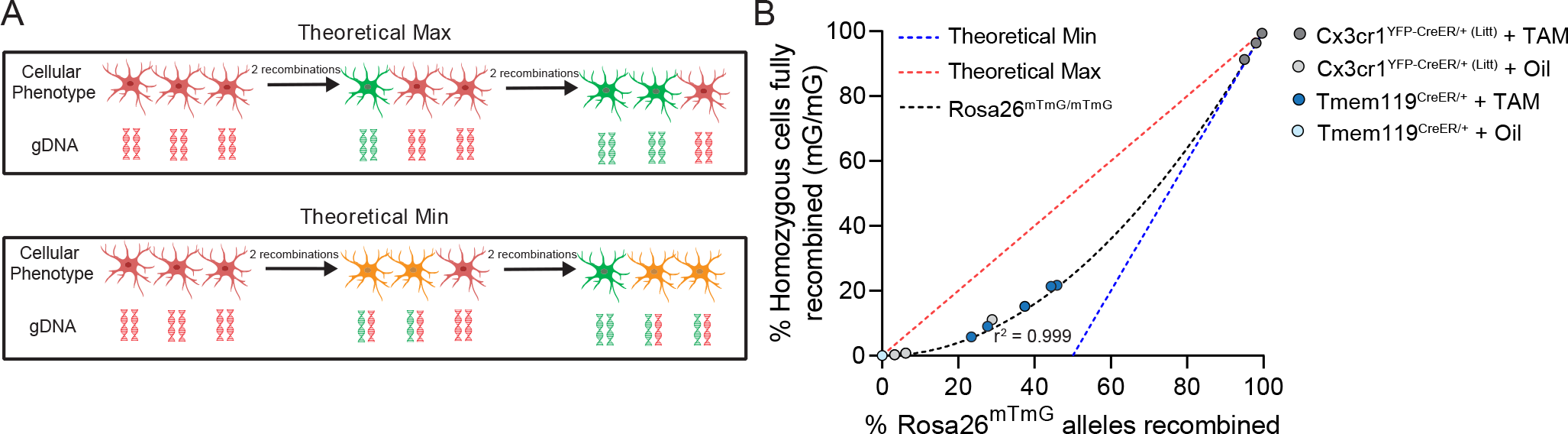
Independent recombination of *Rosa26^mTmG^* alleles in homozygous cells. Related to Figure 4 **A** Diagram of the number of non-recombined (red), singly recombined (orange), and doubly recombined cells (green) obtained after recombination of genomic DNA (gDNA). Diagrams show the theoretical maximum (max; top) and theoretical minimum (min; bottom) of doubly recombined cells for a given number of recombined alleles of gDNA. **B** Plot of the percentage of alleles recombined vs. the number of homozygous cells fully recombined for the theoretical min (blue dashed line), theoretical max (red dashed line), and observed data of *Rosa26^mTmG/mTmG^* recombination in oil-treated *Tmem119^CreER/+^* mice (light blue dots), tamoxifen (TAM)-treated *Tmem119^CreER/+^* mice (dark blue dots), oil-treated *Cx3cr1^YFP-CreER/+^* ^(Litt)^ mice (light grey dots), and TAM-treated *Cx3cr1^YFP-CreER/+^* ^(Litt)^ mice (dark grey dots). The observed data from *Rosa26^mTmG/mTmG^* mice fit to a second-order polynomial curve (black dashed line; *y* = *x^2^*; *r^2^* = 0.999).

**Table S1.**
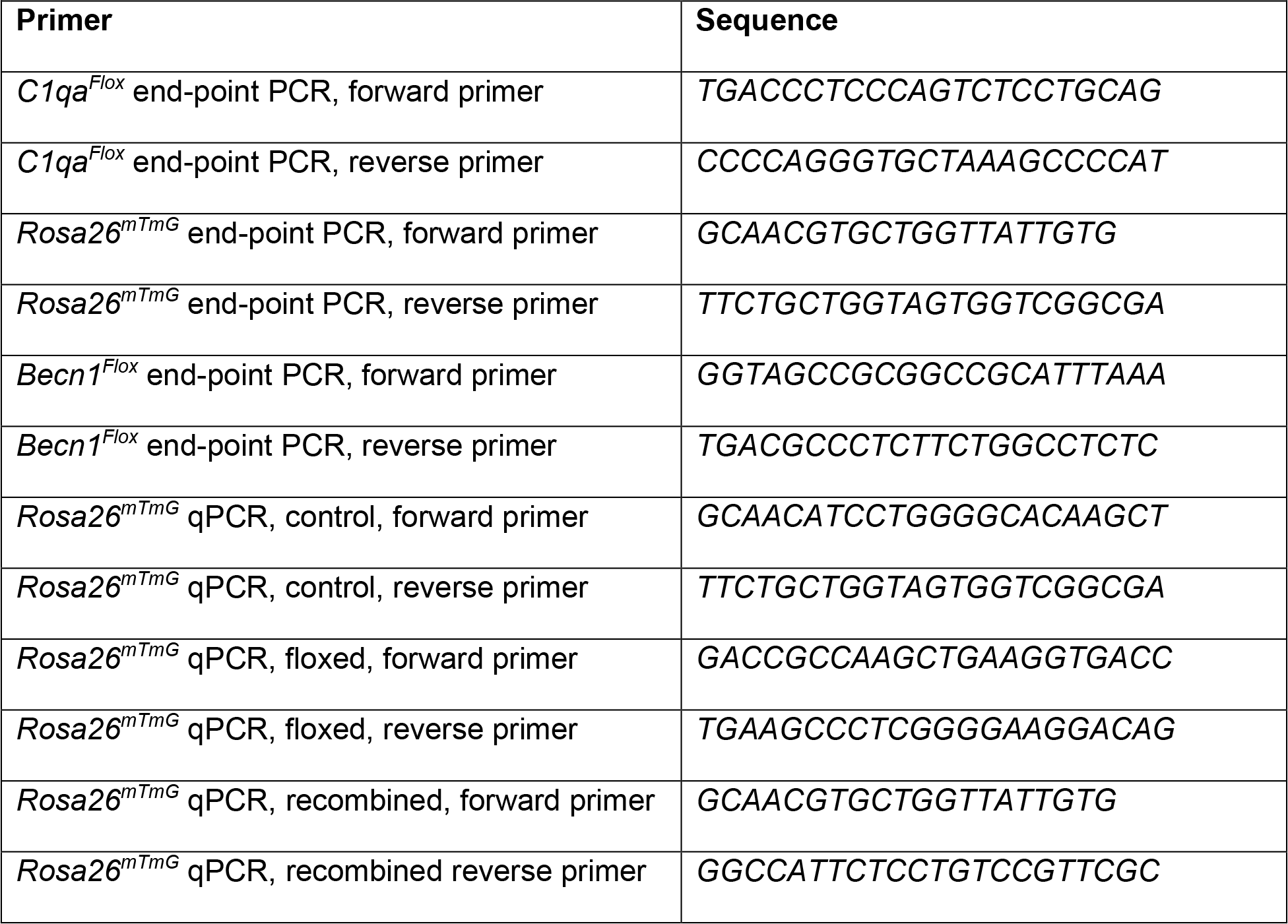
DNA Primers.

